# CRISPR/FnCas12a-mediated efficient multiplex and iterative genome editing in bacterial plant pathogens without donor DNA templates

**DOI:** 10.1101/2022.10.31.514474

**Authors:** Fang Yan, Jingwen Wang, Sujie Zhang, Zhenwan Lu, Shaofang Li, Zhiyuan Ji, Congfeng Song, Gongyou Chen, Jin Xu, Jie Feng, Xueping Zhou, Huanbin Zhou

## Abstract

CRISPR-based genome editing technology is revolutionizing prokaryotic research, but it has been rarely studied in bacterial plant pathogens. Here, we have developed a targeted genome editing method with no requirement of donor templates for convenient and efficient gene knockout in *Xanthomonas oryzae* pv. *oryzae* (*Xoo*), one of the most important bacterial pathogens on rice, by employing the heterogenous CRISPR/Cas12a from *Francisella novicida* and NHEJ proteins from *Mycobacterium tuberculosis.* FnCas12a nuclease generated both small and large DNA deletions at the target sites as well as it enabled multiplex genome editing, gene cluster deletion and plasmid cure in the *Xoo* PXO99^A^ strain. Accordingly, a non-TAL effector-free polymutant strain PXO99^A^D25E, which lacks all 25 *Xop* genes involved in *Xoo* pathogenesis, has been engineered through iterative genome editing. Whole-genome sequencing analysis indicated that FnCas12a did not have a noticeable off-target effect. In addition, we revealed that these strategies are also suitable for targeted genome editing in another bacterial plant pathogen *Pseudomonas syringae* pv. *tomato* (*Pst*). We believe that our bacterial genome editing method will greatly expand the CRISPR study on microorganisms and advance our understanding of the physiology and pathogenesis of *Xoo*.

## Author Summary

Genetic manipulation of bacterial genomes has greatly advanced our understanding of bacterial physiology, which helps us build up cell factories for the production of natural products and breed custom-designed crops for improved disease resistance. Compared with the donor-DNA-mediated homologous recombination methods commonly used in bacterial genome manipulation so far, the donor-DNA-free and CRISPR/Cas-based approaches widely developed in eukaryotic genome editing are more attractive due to its simplicity, versatility, high specificity, and efficiency. However, ectopic expression of Cas proteins causes cell death in many bacteria species, and CRISPR technology has been rarely studied in plant bacterial pathogens. In this work, by co-expressing *mtLigD* and *mtKu* genes, we confirm that the CRISPR/FnCas12a-induced lethal DSBs can be repaired in an error-prone manner, which results in efficient targeted genome editing in both *Xoo* and *Pst* strains. As a result, we have fulfilled various genome editing applications through simple gRNA design and construction, including sequentially knocking out all 25 *Xop* genes in *Xoo* PXO99^A^ strain, plasmid curing, *etc*. This study highlights the importance and promising application of utilizing heterologous NHEJ proteins in genome editing in microorganisms lacking NHEJ machines in the future.

## Introduction

*Xanthomonas* species, rod-shaped gram-negative bacteria, are well known for its pathogenicity in a wide variety of different plant hosts [1]. Among them, *Xoo* causes bacterial leaf blight (BLB) in rice, which is one of the most severe and prevalent rice diseases across the world and result in substantial yield loss during an epidemic. Dissecting the molecular mechanisms of rice-*Xanthomonas* interactions provides insights helpful in developing more effective rice cultivars with durable and broad-spectrum resistance to BLB. In the last decade, clustered regularly interspaced short palindromic repeats (CRISPR)/CRISPR-associated protein (Cas)-mediated genome editing technology, derived from many bacteria and most archaea, has revolutionized everything from basic science to translational application related to microorganisms, plants, animals, and human being [2–7]. However, the application of CRISPR technology in *Xanthomonas* species, even in plant pathogenic bacteria, has been rarely studied.

Our current knowledge about CRISPR-mediated genome editing in bacteria mainly come from a number of human pathogens as well as industrial and laboratory strains, such as *Escherichia coli* [8–10], *Pseudomonas* [11, 12], *Mycobacterium* [13], *Bacillus* [14, 15], *Corynebacterium* [16], *Clostridium* [17], *Streptomyces* [18], *etc*. Generally speaking, the type II CRISPR/Cas9 system[19] is the most widely-used heterologous genome editor in various bacterial strains due to its robust nuclease activity and broad compatibility with other functional proteins (i.e. nucleotide deaminases, transcriptional regulators, DNA polymerase, reverse transcriptase *etc*.), which facilitates DNA fragment deletion and replacement, nucleotide changes, transcriptional depression and activation of target genes as well as artificial gene evolution [9, 20–22]. By contrast, the type V CRISPR/Cas12a system, characterized by a single crRNA and a thymine-rich protospacer adjacent motif (PAM), has been attractive and increasingly applied in bacterial genome editing in recent years [19, 21, 23]. Alternatively, the endogenous type I CRISPR systems, comprising the Cas3 nuclease and multiple effector proteins, have also been successfully reprogrammed to self-edit specific genome regions in its native hosts [24–26].

As reported in prokaryotes so far, double-strand DNA breaks (DSBs) in genomic DNA could be repaired through three different mechanisms: homology-directed repair (HDR), classical non-homologous end joining (NHEJ) and alternative end joining (AEJ; also referred to as microhomology-mediated repair, MMEJ), the latter two mechanisms are independent of DNA repair template [27–29]. However, most bacteria do not have NHEJ/AEJ pathways with a few exceptions, such as *E. coli* [28], *P. aeruginosa* [30], *B. subtilis* [27], *M. tuberculosis* [13], *R. capsulatus* [31], *A. mediterranei* [32], *etc*. Thus, DSB introduced by Cas nucleases in the bacterial chromosome is severely toxic to cells and HDR-mediated genome editing with exogenous donor have been widely utilized in bacteria. Accordingly, multiplex genome editing and genome-scale knockout screening are still challenging in bacteria.

The type I-C CRISPR system has been found in genome-sequenced *Xoo* strains worldwide [33, 34]. It has been characterized by a set of conserved *Cas* genes and a highly variable CRISPR array. Detailed investigation showed that hijacking the endogenous CRISPR system with artificial mini-CRISPR array resulted in self-target killing of the *Xoo* Philippine isolate PXO99^A^ [35]. It was most likely due to unsuccessful DNA repair of Cas3-induced DSBs in PXO99^A^. Indeed, recent study showed that DSBs caused by *Xoo* CRISPR system could be efficiently repaired by heterologous recombinase (λ-Red) with donor templates, allowing precise genome editing in *Xoo* [36]. Here, to develop an easy-to-use versatile genome editing platform in plant bacterial pathogens, we have comprehensively explored the potentials of the CRISPR/Cas12a system from *F. novicida* (FnCas12a) and the crucial NHEJ proteins from *M. tuberculosis* (mtNHEJ) in PXO99^A^. Our data indicates that harnessing both CRISPR/FnCas12a and mtNHEJ achieves highly efficient single and multiple gene knockout, genomic DNA fragment deletion as well as iterative rounds of gene targeting. Also, we have validated that our method is suitable for genetic manipulations in another important plant bacterial pathogen *Pst* DC3000 as well.

## Results

### Development of a CRISPR/FnCas12a-based genome editing system in *Xoo* with heterogenous NHEJ proteins

To develop a versatile bacterial genome editing system, an intermediate CRISPR construct (termed pHZB3) has been designed and generated using exogenous CRISPR/FnCas12a system instead of the endogenous Cas3 nuclease, in which *FnCas12a* was under the control of the inducible tetracycline promoter to reduce its potential toxicity in bacterial cells and the crRNA was directed by the synthetic constitutive j23119 promoter (Fig 1A). Thus, the 23-nt crRNA protospacers towards TTTV PAM for gene targeting can be easily cloned, and the resulting FnCas12a/crRNA-expressing cassettes can be simply released through *Bam*HI digestion and integrated into broad-host-range vectors (i.e. pHM1, pBBR1, *etc*) for the application in diverse bacteria if wanted. With this strategy, the pHM1B3 construct which carried a gRNA targeting the virulence type III secreted effector (T3SE)-encoding gene *XopN* was electroporated into the *Xoo* strain PXO99^A^ (Fig 1B). As shown in Fig. 1C, XopN-crRNA yielded a few colonies on agar plates complemented with the transcriptional inducer anhydrotetracycline (aTc). The significant reduction in transformation efficiency of XopN-crRNA compared to that of empty control pHM1B3 indicated that FnCas12a effectively cleaved the bacterial chromosome and the DSB was toxic to the host cells. Twenty-two survival colonies were randomly picked for genotyping and all were confirmed as wild type (Data not shown), suggesting that endogenous NHEJ and AEJ might not exist or are not efficient enough for DSB repairing in PXO99^A^.

**Fig 1.**
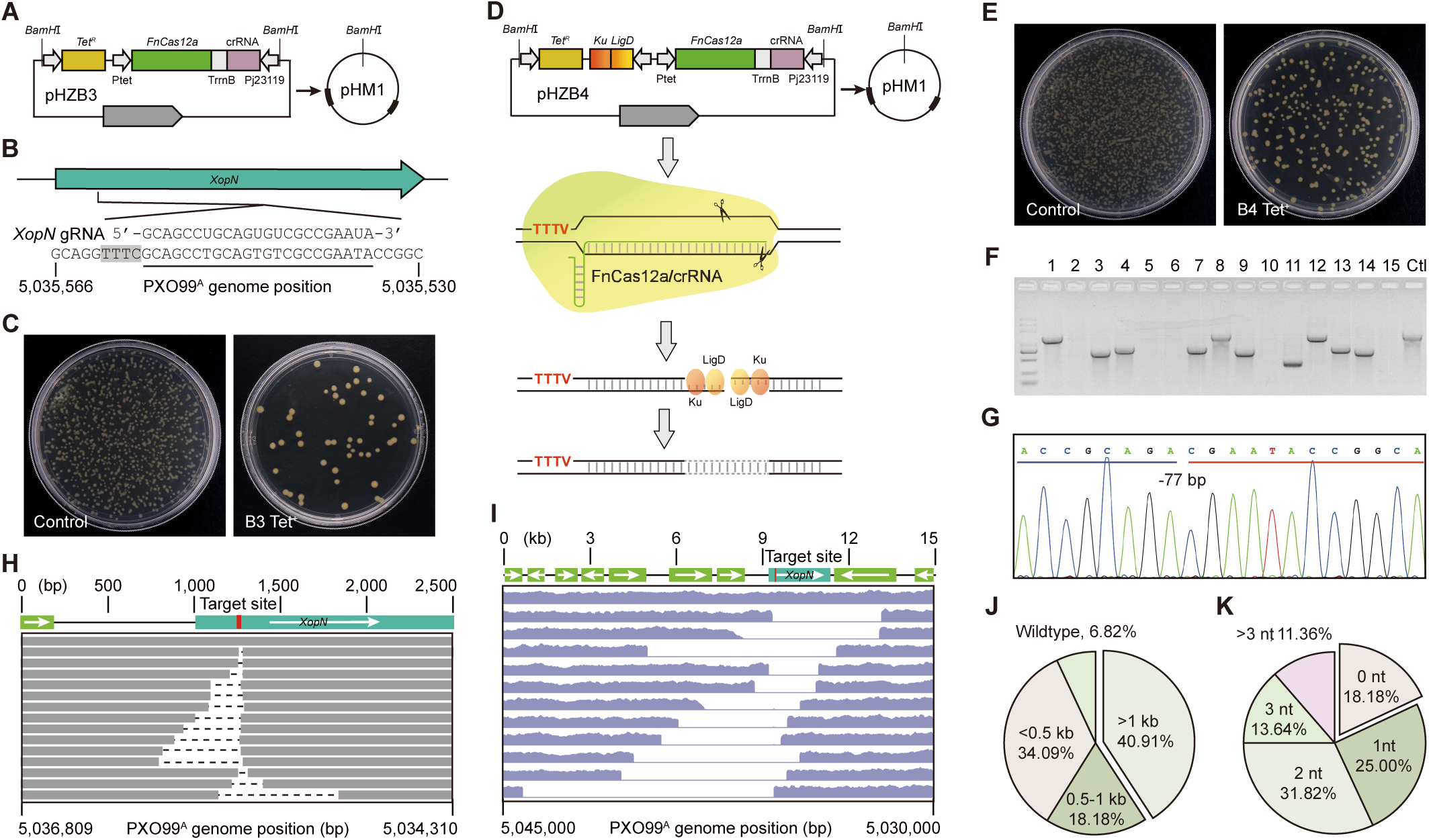
CRISPR/FnCas12a-mediated genome editing in *Xoo* PXO99^A^. **(A)** Schematic illustration of pHM1B3 CRISPR cassette construction. The intermediate CRISPR construct pHZB3 is released through *BamH*I digestion and integrated into the vector pHM1. *Tet^R^*, tetracycline resistance gene; Ptet, tetracy-cline-inducible promoter; FnCas12a, the CRISPR/Cas12a system from *F. novicida*; TrrnB, *E. coli* ribosom-al RNA manipulator rrnB; crRNA, a matured CRISPR RNA; Pj23119, a synthetic constitutive expression promoter; pHM1, a broad-host-range vector. **(B)** The target site of *XopN*. The target region in PXO99^A^ genome is underlined and the PAM sequence is marked by the black shadow. **(C)** Electroporated PXO99^A^ on agar plates. Under the same conditions, except that the control (left plate) didn’t contain protospacer of *XopN*. **(D)** Schematic illustration of pHM1B4 CRISPR cassette construction. It’s updated from **(A)** with a single operon encoding mtLigD and mtKu proteins. These two proteins act as non-homologous end joining (NHEJ) in *Xoo* cells. **(E)** As for **(C)**, but showing the genome editing system of pHM1B4. **(F)** Single colonies in **(E)** were randomly selected for preliminary identification by PCR amplification of a 1.3-kb genomic fragment flanking the *XopN* gene. Ctl, wild-type control. **(G)** Representative Sanger sequencing chromatogram of the deletion mutant. -77 bp, 77-bp deletion. **(H)** Deletion size distribution of independent mutant strains was determined by tiling PCR and Sanger sequencing. Each line represents an independent mutant, the dash lines indicate the deleted region in each mutant, and the target site location is marked in red. **(I)** As for **(H)**, but determined by the whole-genome sequencing (WGS) analysis of 11 PXO99^A^ mutants. **(J)** The pie chart represents the proportion of different bidirectional deletion ranges. (n = 44 individual colonies randomly selected from **(E)**). **(K)** As for **(J)**, but depicting the ratio of the different flanking micro-homology regions used for DSB repair.

Previous studies showed that mtLigD and mtKu, two crucial NHEJ proteins which have been identified in *M. tuberculosis*, also function in DNA end-joining in other bacteria, such as *E. coli* and *Corynebacterium glutamicum* [37, 38]. We assumed that heterogeneous mtNHEJ proteins would repair FnCas12a-induced DSBs in *Xoo* cells. Therefore, pHZB3 has been upgraded with a single operon of *mtLigD* and *mtKu*, resulting in pHZB4 (Fig 1D). The XopN-crRNA was tested again under the same conditions with the pHZB4 strategy. As expected, pHM1B4 showed greatly increased (∼5-fold) transformation efficiency compared to pHM1B3 in repeated experiments (Fig 1E). Forty-four colonies were randomly picked for colony PCR, it revealed distinct patterns of genomic deletions at the target site (Fig 1F). Next, tiling PCR and Sanger sequencing were used to determine the flanking sequences of small deletions (Fig 1G and 1H). While for those (11 colonies) failed in PCR amplification, whole-genome sequencing (WGS) analysis was performed (Fig 1I). As a result, 41 out of 44 colonies (93.18% editing efficiency) were confirmed with bidirectional deletions ranging in size from 23 bp to 8,808 bp, suggesting that DSBs induced by FnCas12 nuclease are efficiently repaired by the ectopically expressed mtNHEJ proteins. Of note, it was observed that 15 colonies (34.09% ratio) carried small deletions less than 0.5 kb and 18 (40.91% ratio) had fragment deletions larger than 1 kb in size (Fig 1J). In addition, 36 colonies (81.82% ratio) were characterized by flanking microhomology regions ranging from 1 to 7 bp at the breakpoint junctions (Fig 1K), implying that both NHEJ and AEJ pathways are responsible for DSB repairing in the presence of mtLigD and mtKu in PXO99^A^.

Another virulence T3SE gene, *XopV*, was further targeted using the pHZB4 strategy (S1A Fig). Colony PCR and Sanger sequencing analysis showed that XopV-crRNA achieved comparable transformation efficiency and generated deletions of varying sizes at the target site in 93.18% (41 out of 44 colonies) of randomly-selected colonies (S1B-D Fig). Consistent with the finding of XopN-crRNA, XopV-crRNA induced both small (< 0.5 kb, 20.45% ratio) and large (> 1 kb, 54.55% ratio) DNA fragment deletions in PXO99^A^ after DSB repair via both NHEJ and AEJ pathways (S1E and S1F Fig). To be mentioned, the dominant occurrence of large deletions was not well correlated to either the amount or induction time of aTc under our assay conditions (Data not shown). Collectively, these data suggest that the combined use of exogenous CRISPR/FnCas12a system and mtNHEJ proteins enables high-efficiency gene knockout in *Xoo* cells.

### Multiplex genome editing and targeted chromosomal DNA fragment deleting in Xoo

We next investigated whether our method could be employed in multiple gene mutagenesis in *Xoo*. Equal amount of above-mentioned pHM1B4 plasmids carrying XopN-crRNA or XopV-crRNA were electroporated into PXO99^A^ cells simultaneously (Fig 2A). Only single editing event (small and large deletions) at either *XopN* or *XopV* target sites, but not co-editing events, was detected after transformation (Fig 2B and 2C), suggesting that multiple gRNAs might need to be constructed in one single plasmid for multiplex genome editing.

**Fig 2.**
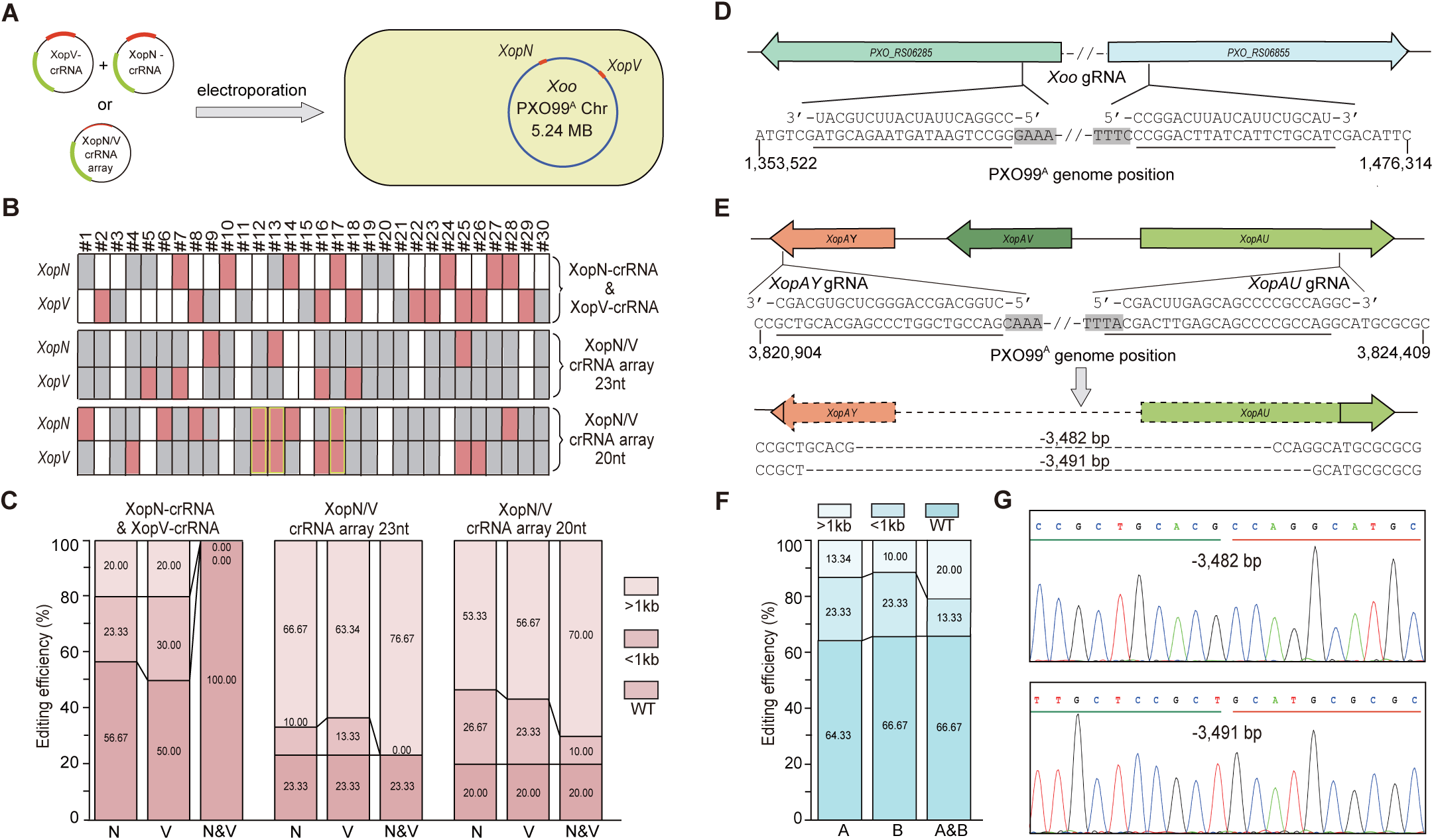
Multiplex gene knockout and accurate chromosomal DNA fragment deletion in *Xoo* **PXO99^A^ by CRISPR/FnCas12a**. **(A)** Schematic illustration of pBBR1B4-mediated simultaneous gene editing of *XopN* and *XopV* in PXO99^A^ with different approaches. Approach I: Equal amounts of plasmid DNA of pHM1B4-XopN-crRNA and pHM1B4-XopV-crRNA were simultaneously electroporated into PXO99^A^. Approach II: A single pHM1B4 plasmid carrying a crRNA array of 20- or 23-nt protospacers was electropo-rated into PXO99A. The relative positions of *XopN* and *XopV* genes in the PXO99^A^ genome (NC_010717.2) are marked in red. **(B)** Thirty independent colonies were randomly picked for genotyping by tilling PCR amplification across ∼3 kb region around the target sites. Each column represents the identities of *XopN* and *XopV* in a single independent colony. The blank, grey, and red boxes denote wild type, large deletions (>1 kb), and small deletions (<1 kb), respectively. Colonies bearing small deletions in both *XopN* and *XopV* are framed by yellow rectangles. #1-30, colony No.; crRNA-XopN & crRNA-XopV, *Xoo* cell transformation with two pHM1B4 plasmid (Approach I); XopN/V crRNA array, *Xoo* cell transformation with a single pHM1B4 plasmid carrying 20- or 23-nt protospacer (Approach II). **(C)** Statistical graphics showing the percentage of various deletion mutations in *XopN* and *XopV* in regard to different genome editing approaches. N, *XopN*; V, *XopV*; N&V, *XopN* and *XopV*. **(D)** Genome editing of two homologous genes (*PX-O_RS06285 and PXO_RS06855*) using crRNA with a 20-nt protospacer. **(E)** Schematic illustration of precise deletion of a 3.84-kb *XopAY/AV/AU* gene cluster using 23-nt paired gRNAs. Survival colonies were genotyped by PCR amplification and Sanger sequencing. The sequences below are the junction sequences with the deletions (dashed lines). The target regions are underlined and the PAM sequences are marked by the black shadow in **(D)** and **(E)**. **(F)** Statistical graphics showing the percentage of various deletion mutations in *PXO_RS06285* and *PXO_RS0685585* using approach II (20-nt protospacer). A, *PXO_RS06285*; B, *PXO_RS06855*; A&B, both *PXO_RS06285*and *PXO_RS06855*. **(G)** Sanger sequencing chromatograms of the deletion mutants shown in **(E)**. The junction sequences are underlined in different colors (green and red).

It’s well known that FnCas12a is capable of processing its own pre-crRNA into mature crRNAs [23], rendering it a versatile tool for multiplex genome editing in bacteria with a single customized CRISPR array consisting of tandem repeats of 19-nt direct repeat and 23-nt protospacer. Therefore, the 23-nt XopN-crRNA and XopV-crRNA were assembled into a crRNA array and electroporated into PXO99^A^ cells with the pHZB4 strategy. We observed that dual gRNAs caused a dramatic decrease in the number of survival colonies in repeated experiments, which were likely resulted from untimely DSB repair and cell death triggered by large chromosomal excision between *XopN* and *XopV* genes. A total number of 30 colonies was genotyped by PCR amplification of the target regions, 23 colonies (76.67% efficiency) were identified with deletion mutations (Fig 2B and 2C). Of note, co-editing of both *XopN* and *XopV* occurred in all positive colonies. However, large DNA deletions (PCR amplification failure across appropriate 3-kb genomic region), instead of small deletions, were observed at all target sites (Fig 2B and 2C). To address this problem, the original 23-nt protospacers were shorten to 20-nt long in order to attenuate the nuclease activity of FnCas12a/crRNA complex in considering of timely DSB repair in transformants. The resulting crRNA array was tested under the same conditions as described above. We observed that 24 out of 30 randomly-selected colonies (80.00% efficiency) had deletions, 3 of which contained small deletions ranging from 9 to 906 bp in both *XopN* and *XopV* (Fig 2B and 2C).

Building on this finding, another 20-nt gRNA was designed to target a conserved region of two homologous genes, *PXO_RS06285* and *PXO_RS06855* which share 91.83% sequence identity in PXO99^A^, and further examined in multiplex genome editing (Fig 2D). Out of 30 randomly-selected colonies, 10 positive colonies (33.33% efficiency) were identified with double gene mutation. Among them, 4 colonies were characterized by small deletions (19-569 bp) in both target genes (Fig 2F). Collectively, these results suggest that the size of deletion mutation in bacterial genome editing could be affected by the activity of CRISPR/FnCas12a system and 20-nt protospacers are likely more suitable for multiplex genome editing in *Xoo*.

Meanwhile, two 23-nt specific gRNAs (XopAY-crRNA and XopAU-crRNA) were designed to target both ends of a 3.84-kb gene cluster, which encode XopAY, XopAV, and XopAU effectors (Fig 2E), and used to investigate if the simultaneous cleavages of chromosome allow the precise deletion of genomic DNA fragment in PXO99^A^. After electroporation, 19 out of 22 randomly-selected colonies (86.36% efficiency) were identified with target deletions. Among them, 2 colonies were characterized by the accurate excision of 3,482-bp and 3,491-bp fragment, which were derived from the direct end-joining between the two predicted cleavage sites of FnCas12a in *XopAY* and *XopAU* (Fig 2E and 2G).

### Construction of PXO99^A^ ploymutant strain lacking all 25 non-TAL effectors

Next, we sought to develop a protocol for iterative rounds of genome editing in PXO99^A^, which allows for unlimited stacking of targeted gene knockouts. In this regard, previously introduced pHM1B4 plasmid must be eliminated prior to the next round of genome editing. To address this question, a crRNA array containing 2 gRNAs, which individually targeted the oriV replicon of pHM1 and the *mtLigD* gene in pHM1B4, respectively, was designed (S2A Fig) and integrated into pHM1B3, resulting in pHM1B3free. Theoretically, both plasmid-curing of any pHM1B4 construct and self-curing of pHM1B3free can be simultaneously achieved through one-step transformation of pHM1B3free due to the inefficient rejoining of linearized plasmids with the disrupted mtLigD protein and the inability of plasmid propagation with disrupted oriV replicon. pHM1B3free was electroporated into the *XopN*-edited PXO99^A^ strain harboring the pHM1B4-XopN-crRNA plasmid mentioned above and the dynamics of plasmid-curing was investigated with different dosages of aTc at different time points. Spectinomycin sensitivity assay and PCR genotyping indicated that 70-75% colonies were plasmid-free after 4 to 12-hour induction of the curing system (S2B-D Fig). Thus, the application of pHM1B3 plasmid enables fast and highly efficient plasmid curing, and the repeated use of pHM1B4-crRNA and pHM1B3free constructs allows us to establish a convenient workflow for *Xoo* strain construction (Fig 3A).

**Fig 3.**
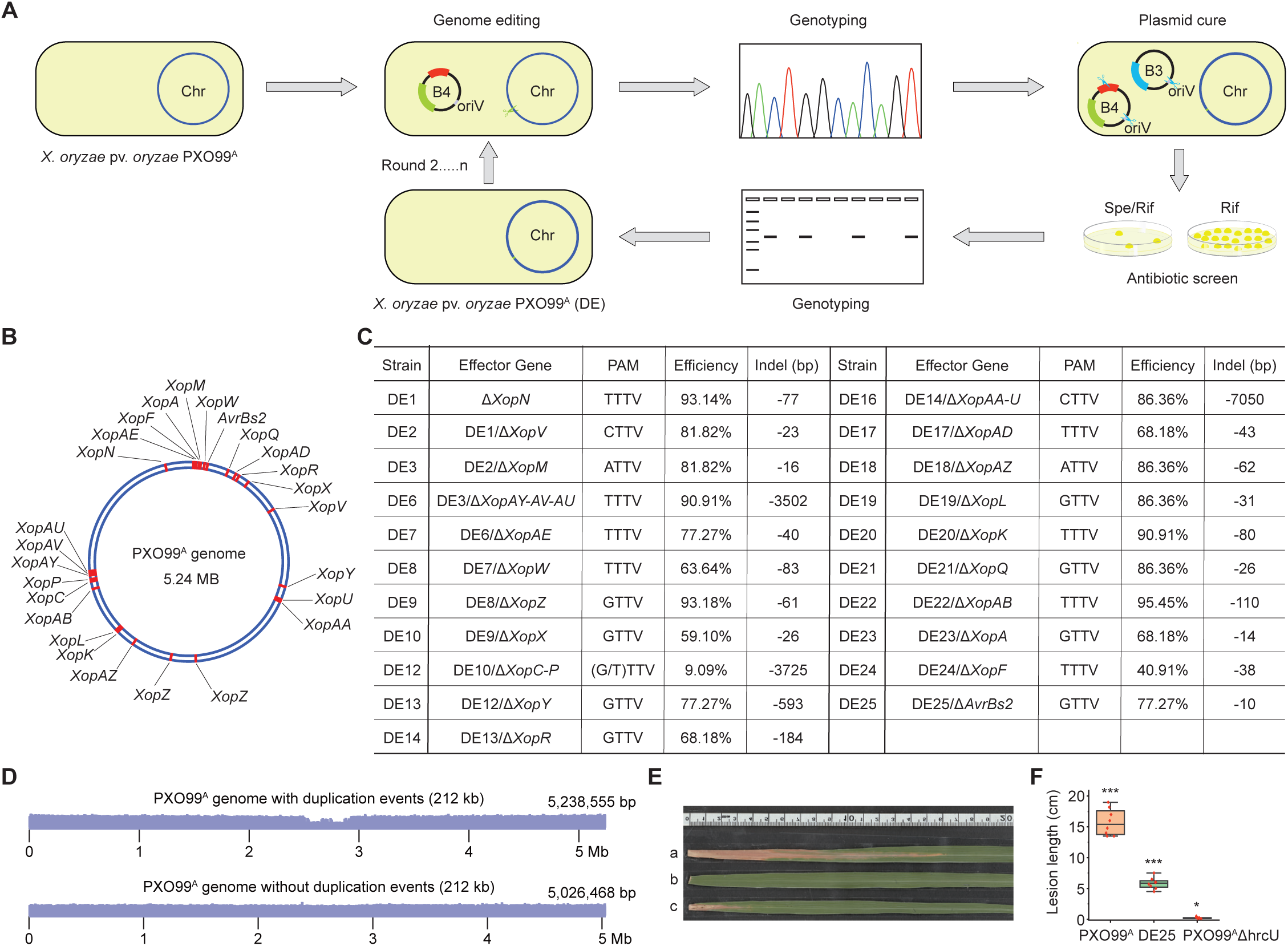
Iteratively knocking out all 25 non-TAL effector genes in Xoo PXO99^A^. **(A)** A protocol for iterative rounds of genome editing in PXO99^A^. The pHM1B4 plasmid was transformed into PXO99^A^ competent cells by electroporation. Survival colonies were genotyped by Sanger sequencing of the targeted gene, and then the mutant strains that meet the requirements were selected for plasmid-curing. Determination of successful plasmid elimination was done by screening in media and PCR assay, and a plasmid-free mutant was used for the next round of knockout operations. Spe, spectinomycin; rif, Rifampicin. **(B)** The positions of all 25 non-TAL effectors in the PXO99^A^ genome (NC_010717.2) are marked in red. **(C)** The table shows the name, target gene(s), PAM sequence, efficiency and deletion size of the Xop mutants obtained in each round of genome editing. **(D)** Genome-wide coverage of PXO99^A^ when aligned back to the genome with the 212 kb duplication event (top) and the genome without the 212 kb duplication event (bottom). **(E)** Disease phenotypes of Kitaake after inoculation with *Xoo* strains. a, PXO99^A^; b, PXO99^A^ΔhrcU; c, PXO99^A^DE25. **(F)** Box plots of mean disease lesion lengths (cm) on Kitaake. Lesions were measured 14 days after inoculation. The red scatters indicate five repetitions. Center lines show medians; box limits indicate the 25^th^ and 75^th^ percentiles determined in the R software package; whiskers extend 1.5 times the inter-quartile range from the 25^th^ and 75^th^ percentiles. Student’s t-test: *P < 0.05, ***P < 0.001.

The highly virulent *Xoo* strain PXO99^A^ encodes 18 transcription activator-like effectors (TALEs) and 25 non-TAL effectors (Xops). Of which a number of Xop proteins (i.e. XopR, XopY, XopP, *etc*.) have been already confirmed as virulence factors in suppressing immunity to *Xoo* in rice [39–41]. To provide new research tool for better probing rice-*Xoo* interactions, we aimed to generate a disarmed PXO99^A^ polymutant strain lacking all Xop effectors through iterative genome editing. Thus, the remaining 24 annotated *Xop* genes in PXO99^A^ΔXopN strain were knocked out consecutively to produce PXO99^A^D25E strain with small deletions exclusively residing in the coding region of each *Xop* gene (Fig 3B). An overview of genome editing of each *Xop* and effector gene cluster is provided in Figs 3C, S3 and S4. In summary, pHM1B4 resulted in high editing frequencies ranging from 40.91 to 95.45% when single gRNA being used to target each *Xop* gene in each round (Fig 3C). As for the other *Xops* arranged in gene clusters, the crRNA arrays achieved DNA fragment deletion efficiency of 90.91%, 9.09%, and 86.36% for *XopAY-AV-AU*, *XopC-P*, and *XopAA-U*, respectively (Fig 3C). Notably, crRNAs towards four types of NTTV PAMs (ATTV, CTTV, TTTV, and GTTV) were included in our experiment, its high editing efficiency indicates that, different to other bacteria, CRISPR/FnCas12a recognizes TTV PAM in *Xoo* cells. Overall, our iterative genome editing system facilitates *Xoo* strain construction with simple gRNA manipulations compared to homologous donor-mediated traditional methods.

Furthermore, the PXO99^A^D25E strain was subjected to WGS analysis to evaluate the genome-wide off-target DNA mutations induced by FnCas12a nuclease. Besides the identified on-target DNA deletions in 25 *Xop* genes, 3 single nucleotide polymorphisms (SNPs) and two small indels (1-nt deletion and 7-nt insertion) were detected in the PXO99^A^D25E genome (S5 Fig). Among these off-target sites, only 1 SNP was located in the coding region, resulting in the substitution of asparagine with aspartic acid (N2D) of RS25085 (S5D Fig). Given the occurrence of 5 crRNA-independent mutations (likely to be spontaneous mutations) in 42 times of FnCas12a-mediated editing, including both gene targeting and plasmid cure, we presume that FnCas12a nuclease with stringently-designed protospacer does not cause obvious off-target mutations in bacterial genome editing. In addition, we verified that there is only one copy of *XopZ* in our PXO99^A^ and PXO99^A^D25E strains (Fig 3D), similar to the report of the PXO99^A^-s reference genome characterized by the 212-kb tandem duplication collapsed [42]. Finally, we inoculated rice variety Kitaake with PXO99^A^D25E, PXO99^A^, and the type III secretion system (T3SS)-deficient PXO99^A^ΔhrcU mutant which is unable to deliver T3SEs. The lesion length caused by PXO99^A^D25E was significantly decreased compared to PXO99^A^, but slightly higher than PXO99^A^ΔhrcU (Fig 3E and 3F), highlighting the critical virulence function of the non-TAL effector repertoire in the pathogenesis of PXO99^A^ on rice.

### Application of CRISPR/FnCas12a in *Pst*

We also investigated pHZB4-mediated genome editing in *Pst*, which is another intensively studied bacterial pathogen in dicot plants but does not bear any native CRISPR/Cas system. A gRNA was designed to target the T3SE gene *HopAB2* residing on the circular chromosome (Fig 4A and 4B). The pHZB4 intermediate carrying HopAB2*-*crRNA was integrated into broad-host-range vectors pBBR1MCS-2 in this case and the resulting pBBR1B4-HopAB2-crRNA construct was electroporated into *Pst* DC3000. As observed, CRISPR/FnCas12a also resulted in decreased transformation efficiency of *Pst* DC3000, which is comparable to PXO99^A^. Twenty-two colonies were randomly picked from 46 colonies for genotyping and 8 colonies (36.37% efficiency) were identified with fragment deletions of various sizes at the target site. Sanger sequencing analysis of small deletions further verified the occurrence of 438 to 714-bp deletions in *HopAB2* (Fig 4C). These data indicate that the combined use of exogenous CRISPR/FnCas12a system and mtNHEJ proteins enables efficient gene knockout in *Pst* as well.

**Fig 4.**
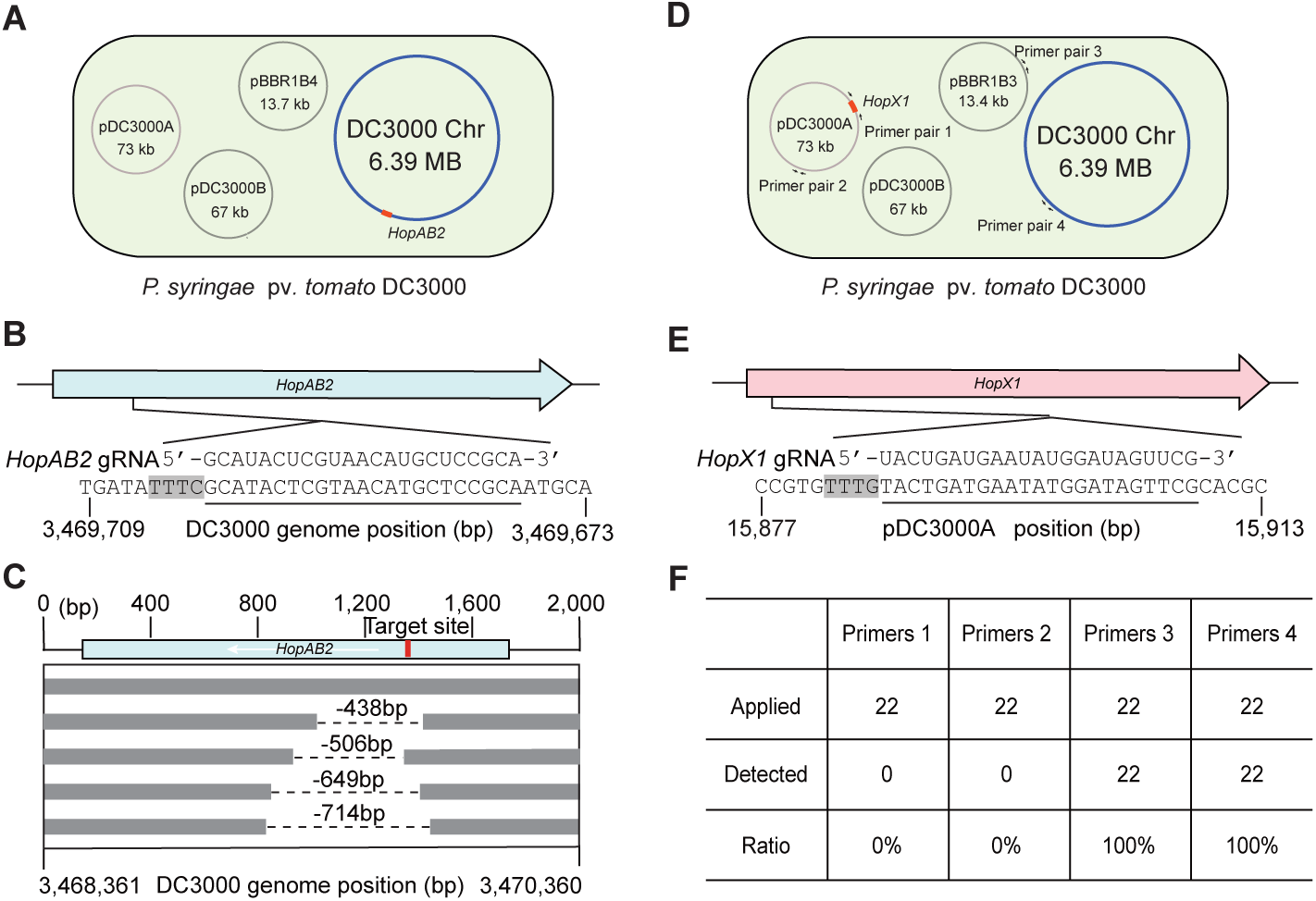
CRISPR/FnCas12a-mediated genome editing in *Pst* DC3000. **(A)** Schematic illustration of pBBR1B4-mediated gene knockout of *HopAB2* residing on the circular chromosome in *Pst* DC3000. The relative position of *HopAB2* on the chromosome is marked in red. **(B)** The target site of *HopAB2*. **(C)** Deletions detected in *HopAB2* knockout mutants by colony PCR and Sanger sequencing. Each line represents an independent mutant, the dash lines indicate the deleted region in each mutant, and the target site location is marked in red. **(D)** Schematic illustration of pBBR1B3-mediated pDC3000A plasmid curing of *Pst* DC3000 by targeting *HopX1*. The relative positions of *HopX1* on pDC3000A and PCR amplification regions are shown in red and by arrows, respectively. The primer sequences are listed in S3 Table. **(E)** The target site of *HopX1*. **(F)** Summary of the plasmid-curing frequency using 22 randomly-selected colonies and primer pairs shown in **(D)**. The target regions are underlined and the PAM sequences are marked by the black shadow in **(B)** and **(E)**, respectively.

Considering that native plasmids in many *Pst* strains are important or essential for host-pathogen interactions and the fast plasmid-curing methods are always attractive to researchers, we next employed a gRNA to target *HopX1*, which encodes another T3SE but presents on the 73-kb native plasmid pDC3000A (Fig 4D and 4E), in an attempt to remove pDC3000A with the pHZB3 strategy. After electroporation, pBBR1B3-HopX1-crRNA yielded a large number of colonies of *Pst* DC3000 on medium plate. Colony PCR was performed using 4 pairs of primers to examine the presence/absence of plasmid pDC3000A and pBBR1B3, and the identity of DC3000 chromosome, respectively, in 22 randomly-selected colonies (Fig 4D). As a result, pDC3000A was eliminated in 100% of the *Pst* cells (Fig 4F), suggesting that CRISPR/FnCas12a nuclease enables high-efficiency curing of native plasmids in *Pst* in one-step transformation.

## Discussion

Plant infection by bacterial pathogens, such as *Xanthomonas*, is complicated and both are in a continuous arms race to battle for survival in nature [1]. The understanding of the bacterial life cycle, pathogenesis and colonization in hosts will help us to breed crops with broad-spectrum resistance as well as to develop bacterial control agents for field application. Also, *Xanthomonas* strains are important resources to produce metabolites for various applications in human society [43]. Therefore, powerful and efficient genetic manipulation methods for *Xanthomonas* are always in high demand. Compared to traditional gene knockout methods based on homologous recombination in *Xanthomonas* [44], CRISPR-mediated genome editing technology is more promising due to its simplicity, target specificity and high efficiency. However, different from the CRISPR craze in eukaryotes [45], the relevant studies in prokaryotes lag far behind, mainly because of high cellular toxicity of CRISPR systems. Cas9 nuclease cause severe cell death in many bacteria, even the catalytically dead Cas9 (dCas9) can produce strong defects or kill *E. coli* by strongly blocking gene expression with specific 5-bp seed sequences at the PAM-proximal region [46]. Therefore, we turned our attention to Cas12 nuclease given its less toxicity and the ability of processing its own pre-crRNA in bacteria.

Three repair mechanisms have been reported for repairing the lethal DSBs in bacteria so far [27–29]. The HDR pathway is prevalent whereas the NHEJ pathway is available in only 20-25% of bacteria [47]. It leads to common use of homologous repair template in CRISPR-based bacterial genome editing, which achieves higher editing efficiency but still require the laborious and time-consuming cloning of homology arms and cannot apply in a large-scale. Therefore, reconstitution of an efficient error-prone NHEJ pathway for repairing FnCas12a-generated DSBs in genome editing is more attractive since its application is convenient and can be scaled up when needed. The NHEJ machine in bacteria is well known that the Ku protein forms a homodimer, binds the initial break and recruits the multifunctional LigD to repair DSBs [37, 47]. Thus, we integrated the heterogenous NHEJ proteins (mtLigD and mtKu) from *M. tuberculosis* with our inducible CRISPR/FnCas12a nuclease system, and successfully achieved highly efficient knockout of the target effector genes in PXO99^A^. Further use of short gRNAs to attenuate DNA targeting by FnCas12a demonstrates that the ectopic expression of both CRISPR and mtNHEJ proteins is also functional in multiplex genome editing. However, we propose that it could be further optimized by enhancing the repair capacity of heterogenous DNA repair machines, such as employing NHEJ proteins from other bacterial species, using codon-optimized NHEJ genes, artificially engineering NHEJ proteins for high-activity variants, *etc*. Anyhow, we assume that our genome editing strategy present in this study is also suitable for developing the relevant tools in other bacterial species, even in other microorganisms.

The pHZB4 strategy dominantly produces DNA fragment deletions (23 bp to 8 kb deletion in *Xoo* and >438 bp deletion in *Pst* in our study) other than insertions at regions of microhomology (1∼7 bp), we reason that the output of genome editing using our method is highly corelated to the distribution of direct microhomologous repeats flanking the cleavage site on the genomic DNA in different bacterial species. Anyhow, the occurrences of both small and large deletions in *Xoo* genome editing allow us to disrupt the target genes precisely without affecting overlapping open reading frames and nearby genes. In addition, the ability of accurate excision of genomic DNA with multiple gRNAs facilitates minimizing the bacterial genome, allowing us to study the pan-genome and evolution of bacterial pathogens.

Given the high toxicity of Cas nucleases, CRISPR interference (CRISPRi)-mediated knockdown approach which uses dCas proteins to temporarily disrupt expression of target genes besides complex Tn-based method have been usually applied in large-scale screen in bacteria [48]. Technically, our high-efficiency pHZB4 strategy opens the prospect of genome-scale mutagenesis in bacteria using Cas nuclease, it provides a mechanistically distinct method from CRISPRi for systematic genetic analysis. Alternatively, one of the important outcomes of our work is the iterative genome editing method with well-designed gRNAs in *Xoo*, it enables the construction of “build-your-own” cell factories for producing metabolites (i.e. xanthan gum, *etc*.) as well as chassis strains (i.e. PXO99^A^D25E, *etc*.) for probing functionally redundant effectors and plant innate immune system.

In conclusion, a donor-free, high-efficiency, targeted gene knockout method has been well established in bacterial plant pathogens by employing both inducible CRISPR/FnCas12a system and heterogenous mtNHEJ proteins in this study. It enables us to conveniently construct bacterial strains regardless of target gene numbers as well as large-scale gene knockout screen regardless of the presence/absence of endogenous end joining repair machines. Our detailed study gives novel insights into the development of targeted genome editing technologies in diverse microorganisms, and undoubtedly accelerates both the fundamental study and applied biological control research in plant pathology as well as metabolic engineering of *Xanthomonas* in the industry.

## Materials and Methods

### Bacterial strains, plasmids and primers

All bacterial strains and plasmids used in the study along with their relevant characteristics and source are described in S4 Table. Primers used for plasmids construction and PCR amplifications are provided in S3 Table.

### Bacterial growth conditions

*X. oryzae* pv. *oryzae* strains were grown in Nutrient Broth (NB, Nutrient Broth, Difco, BD, 8g/L) or Nutrient Broth plate supplemented with 1.5% agar (NA) at 28°C. *P. syringae* pv. *tomato* strains were grown in King’s B (KB) medium (Protease peptone No.3 20g/L, K_2_HPO_4_ 1.5g/L, Glycerol 15g/L, with or without 1.5% agar, add 3.2 mL of 1M MgSO_4_ after autoclaved) at 28℃. *E. coli* strains were grown in Luria Bertani (LB) media with or without 1.5% agar at 37℃. Antibiotics were used at the following final concentrations unless otherwise noted: Ampicillin (Amp), 50 µg/ml; Spectinomycin (Spe), 100 µg/ml; Kanamycin (Kan), 50 µg/ml; Anhydrotetracycline (aTc), 200 µg/ml; Rifampin (Rif), 50 µg/ml.

### Plasmid construction

The 4,394-bp fragment consisting of *FnCas12*-expression cassette and crRNA-expression cassette were codon-optimized for suitable expression in PXO99^A^ and synthesized by Sangon Biotech (Shanghai, China) (S1 Table). The 4,394-bp fragment digested by *Acc*65I/*Spe*I (Thermo Scientific) was cloned into the vector backbone of pENTR4-gRNA4 which was amplified with the primer pair pENTR4-F1/pENTR4-R1 and Phanta Max Super-Fidelity DNA Polymerase (Vazyme, China), resulting in pENTR4-FnCas12a. Tet^R^ (the repressor of the tetracycline resistance element) insertion of was performed by ClonExpress II One Step Cloning Kit (Vazyme, China) with *Spe*I-digested pENTR4-FnCas12a and Tet^R^ amplicon of plasmid pFREE vector (Addgene, #92050) with the primer pair Tet-F3/Tet-R3, resulting in pHZB3. The *mtLigD/mtKu*-expression cassette was codon-optimized for suitable expression in PXO99^A^, synthesized by Sangon Biotech (Shanghai, China) (S1 Table) and then inserted into the pHZB3 through *Spe*I digestion and DNA ligation, resulting in pHZB4.

For the construction of FnCas12a/crRNA plasmids, the 26-nt or 23-nt complementary oligos corresponding to each target site (S3 Table) and carrying appropriate 3-nt adaptor were phosphorylated, annealed, and inserted into the *Sap*I-digested pHZB3 or pHZB4, resulting in pHZB3-crRNA or pHZB4-crRNA, respectively. Similarly, crRNA arrays, composed of tandem repeats of 19-nt direct repeat and 23-nt or 20-nt protospacer, were synthesized by Sangon Biotech (S3 Table) and inserted into the *Sap*I-digested pHZB3 or pHZB4 as described above.

Next, the FnCas12a/crRNA-expression cassettes in pHZB3 and pHZB4 were cloned into pHM1 vector through *Bam*HI digestion and DNA ligation, resulting in pHM1B3-crRNA and pHM1B4-crRNA for genome editing in PXO99^A^ stains, respectively. Similarly, the FnCas12a/crRNA-expression cassettes were cloned into pBBR1MCS-2 to generate pBBR1B3-crRNA and pBBR1B4-crRNA for genome editing in *Pst* DC3000 stains, respectively.

### Competent cell preparation and electroporation

The plasmids used in this study were delivered into *Xoo* PXO99^A^ and *Pst* DC3000 by electroporation. Electro competent cells preparation and electro-transformation were performed as following. 1-mL overnight culture of PXO99^A^ was 1:10 diluted into 10 mL of fresh medium and incubated at 28℃ until OD_600_ reaches 0.7-1.0. The cells were harvested by centrifugation at 5,000g for 10 min. The supernatant was discarded and the cells were washed twice with 10 mL of chilled, sterile MilliQ water at 4℃. Finally, the cells were resuspended in 1 mL (1/10 original volume) of 10% glycerol, and divided into 100 μL aliquot for the subsequent experiments.

The plasmids were transformed into PXO99^A^ by electroporation with the parameters of 2500 V, 2 mm cuvette (Bio-Rad). The mixture was recovered for 2-3 h in 1 mL of NB medium at 28℃ and then cultured for 2 h after adding aTc to a final concentration of 200 μg/mL. The colonies containing pHM1 were selected on NA plates containing 100 μg/mL Spe. Similar procedures were performed to preparation DC3000 electrocompetent cells and transformation using KB medium.

### Plasmid curing

*Xoo* PXO99^A^ strains harboring various pHM1B4-crRNA plasmids were grown in 10 mL of NB complimented with 100 µg/ml Spe to prepare electrocompetent cells. 200 μL of competent cells were transformed with 1μg of pHM1B3free plasmid by electroporation and recovered for 2-3 hours in 2 mL of NB medium at 30℃ with shaking. After that, a gradient of final concentration of aTc (200 ng/mL, 500 ng/mL, 1000 ng/mL, 2000 ng/mL, 3000 ng/mL) was added, and cell cultures at different induction time points (1, 2, 4, 6, 8, 10, and 12 hours) were plated on non-selective NA plates. A number of colonies for each sample were grown on NA plates with/without Spe. Loss of the plasmids was verified by loss of Spe resistance. Plasmid-free colonies were further confirmed by PCR amplification of pHM1 backbone.

### DNA extraction and PCR genotyping

Bacterial genomic DNA was isolated from each sample using the SteadyPure Bacteria Genomic DNA Extraction Kit (Accurate Biotechnology Co., Ltd, China) according to the manufacturer’s protocol. Tilling PCR amplification across the targeted genomic region was carried out with 2×Taq Master Mix (Vazyme, China) with specific primers listed in S3 Table, the size of PCR products is 1-1.5 kb. Shifted gel bands of PCR product was subjected to Sanger sequencing for identifying mutations. As for the colonies with no PCR product, genomic DNAs were subjected to WGS.

### Whole-genome sequencing library construction and data analysis

RNA-free PXO99^A^ genomic DNAs (0.2 μg) were used to construct the DNA libraries using a NEBNext Ultra DNA Library Prep Kit for Illumina (NEB, USA) following the manufacturer’s instructions. DNA libraries were sequenced on the Illumina platform in the 150-nt paired-end mode with around 200X coverage.

The sequencing reads were mapped to PXO99^A^ genome downloaded from NCBI (NC_010717.2) via BWA [49] and sorted using samtools (v1.9) [50] . In addition, the genome without the large 212 kb duplication was created through removing the DNA sequence fragment between 2,502,622-2,714,708 [34]. Genome-wide coverage was calculated using in-house python program. The Genome Analysis Toolkit (GATK v4.2) was used to mark duplicated reads and recalibrate base qualities [51]. To identify high-quality genetic changes at the genomic scale, we applied three independent germline variant-calling methods: GATK, LoFreq [52], and Strelka2 [53]. SNPs and indels identified through all three methods were combined together and shown in IGV browser [54].

### Rice growth, inoculation and statistical analysis

Rice plants of the *Geng* cultivar Kitaake were grown in growth chamber under the following conditions: 30/25℃ (day/night) and 75% relative humidity and photoperiod of 12 h in light/dark. Fully expanded leaves were inoculated with bacterial suspensions (OD_600_ = 0.5) at the four-leaf seedling stage using the leaf-tip-clipping method. Disease symptoms were scored by measuring lesion length 2 weeks post inoculation (DPI). The statistical significance between samples were analyzed by Student’s t-test (p < 0.001).

### Supporting information

**S1 Fig. Targeted gene editing of *XopV* in PXO99^A^ with the pHZB4 strategy.**

**S2 Fig. Fast and efficient curing of exogenous plasmids in PXO99^A^ with the pHZB3 strategy.**

**S3 Fig. Mutation information on *Xop* genes in the PXO99^A^D25E strain generated by iterative genome editing.**

**S4 Fig. Mutation information on *Xop* genes in the PXO99^A^D25E strain generated by iterative genome editing (continued).**

**S5 Fig. Off-target mutations of PXO99^A^ DE25 detected through WGS.**

**S1 Table. The complete nucleotide sequences of the codon-optimized *FnCas12a* and *mtLigD/mtKu* fragments.**

**S2 Table. Target gene list for genome editing in this study.**

**S3 Table. Oligonucleotides used in this study.**

**S4 Table. Bacterial strains and plasmids used in this study.**

## Data Availability Statement

All relevant data are within the manuscript and its Supporting Information files.

## Acknowledgements

The authors thank Dr. Houxiang Kang for help in data analysis, Dr. Bin Ren for assistance in pathogen inoculation experiment, and Dr. Hailei Wei for providing pBBR1MCS-2 plasmid.

## Author contributions

**Conceptualization:** Huanbin Zhou, Xueping Zhou, Jie Feng, Gongyou Chen, Congfeng Song

**Investigation:** Fang Yan, Jingwen Wang, Sujie Zhang, Zhenwan Lu, Shaofang Li

**Resources:** Jin Xu, Gongyou Chen, Congfeng Song, Zhiyuan Ji

**Supervision:** Huanbin Zhou

**Writing-original draft:** Fang Yan, Sujie Zhang, Huanbin Zhou

**Writing-review & editing:** Huanbin Zhou

## Figure Legends

**S1 Fig.**
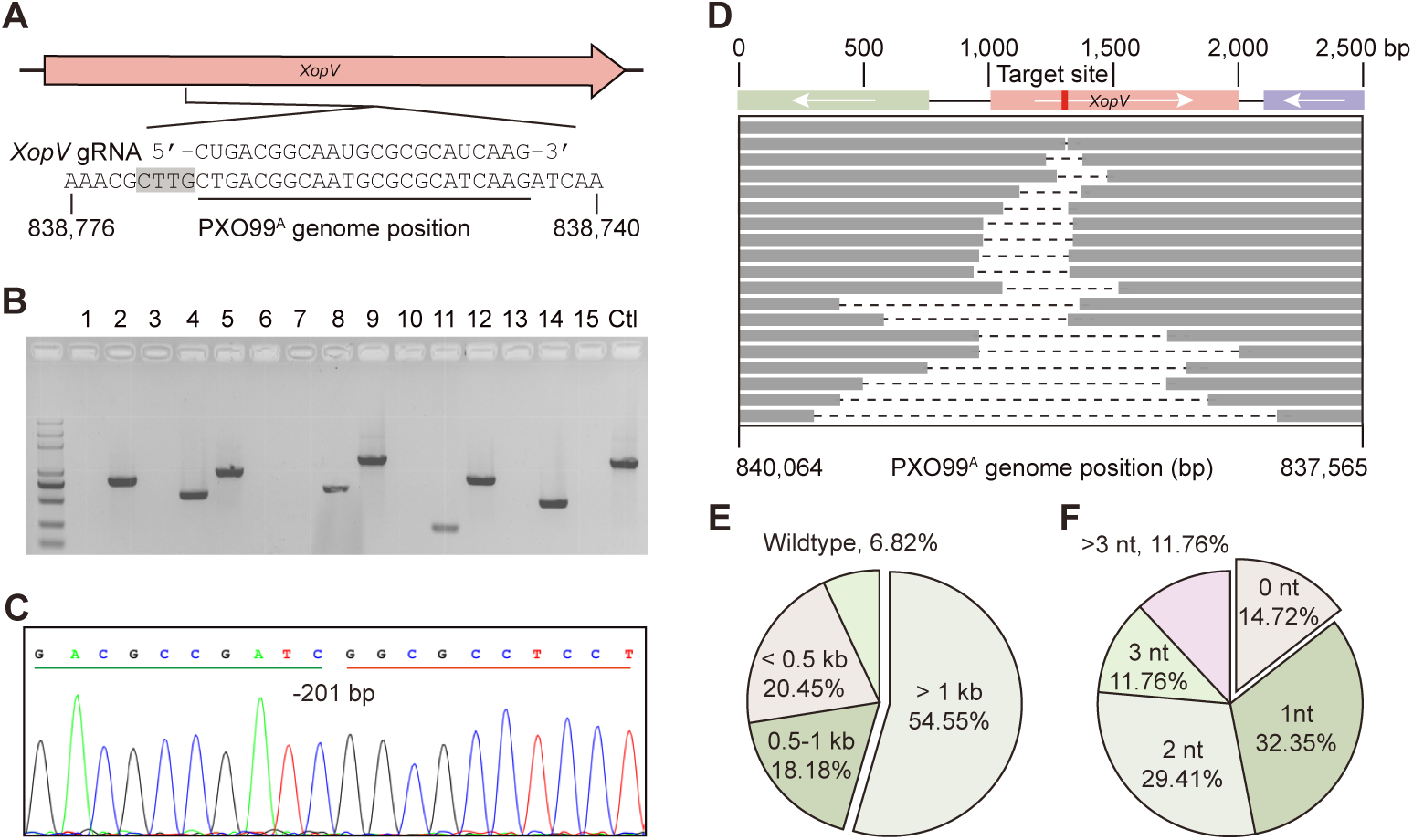
Targeted gene editing of *XopV* in *Xoo* PXO99^A^ with the pHZB4 strategy. **(A)** The target site of *XopV*. The target region in the PXO99^A^ genome is underlined and the PAM sequence is marked by the black shadow. **(B)** Single colonies were randomly selected for preliminary identification by PCR amplification of a 1.1 kb genomic fragment flanking the *XopV* gene. Ctl, control. **(C)** Representative Sanger sequencing chromatogram of the deletion mutant. -201bp, 201-bp deletion. **(D)** Deletion size distribution of mutants determined using tiling PCR and Sanger sequencing. Each line represents an independent mutant, the dash line in the middle indicates the deleted portion of each mutant, and the target site location is marked in red. **(E)** The pie chart represents the proportion of different bidirectional deletion ranges. (n = 44 individual colonies randomly selected). **(F)** As for (E), but depicting the ratio of the different flanking micro-homology regions used for DSB repair.

**S2 Fig.**
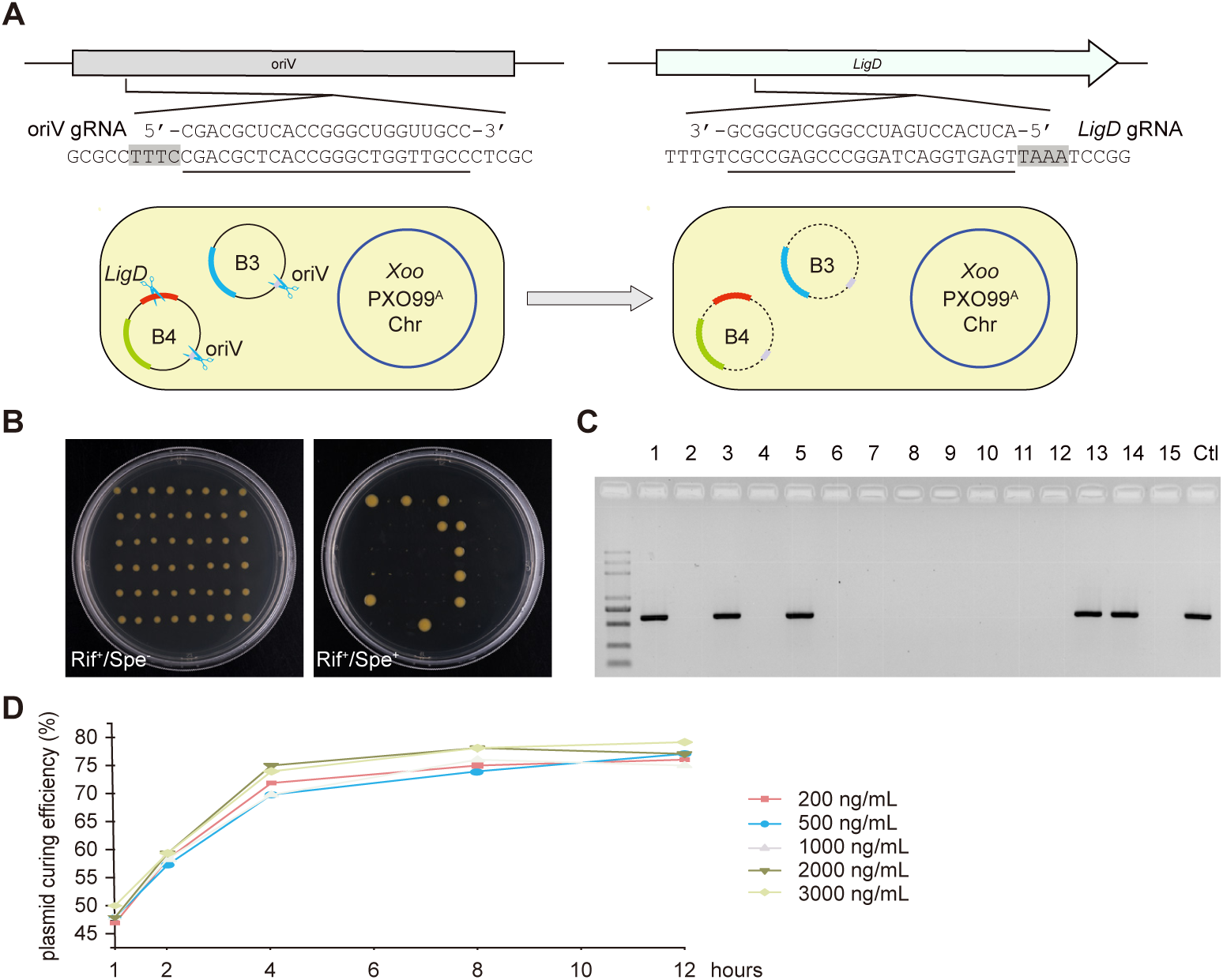
Fast and efficient curing of exogenous plasmids in *Xoo* PXO99^A^ with the pHZB3 strategy. **(A)** CRISPR/FnCas12a nuclease-mediated cleavages of both oriV replicon and *mtLigD* gene resulted in simultaneous removal of pHM1B3 and pHM1B4 plasmids. A crRNA array was designed to target the oriV replicon of pHM1 and the *mtLigD* gene in pHM1B4, respectively. The positions of the target sites in the PXO99^A^ genome are underlined and the PAM sequences are marked by the black shadow. The expression cassettes of *FnCas12a*/XopN-crRNA, *mtKu/mtLigD*, and FnCas12a*/*oriV /mtLigD-crRNA are depicted as yellow-green, red, and deepskyblue stripes. B3, pHM1B3free plasmid; B4, pHM1B4-crRNA plasmid. **(B)** Selection of the plasmid-free PXO99^A^ΔXopN strains on NB plates with/without spectinomycin after pHM1B3free transformation. Spe-, without spectinomycin; Spe+, with spectinomycin added. **(C)** Representative agarose image for the presence/absence of pHM1 plasmid in PXO99^A^ΔXopN. Colonies in (B) were randomly selected for PCR amplification. Ctl, control. **(D)** Time course characterization of the pHM1B3free plasmid-curing system in PXO99^A^ΔXopN cells bearing pHM1B4-XopN-crRNA. The percentage of plasmid-free cells was calculated at induction time 1, 2, 4, 6, 8, 10, 12 hours after treatment of aTc in different dosage (200-3,000 ng/mL).

**S3 Fig.**
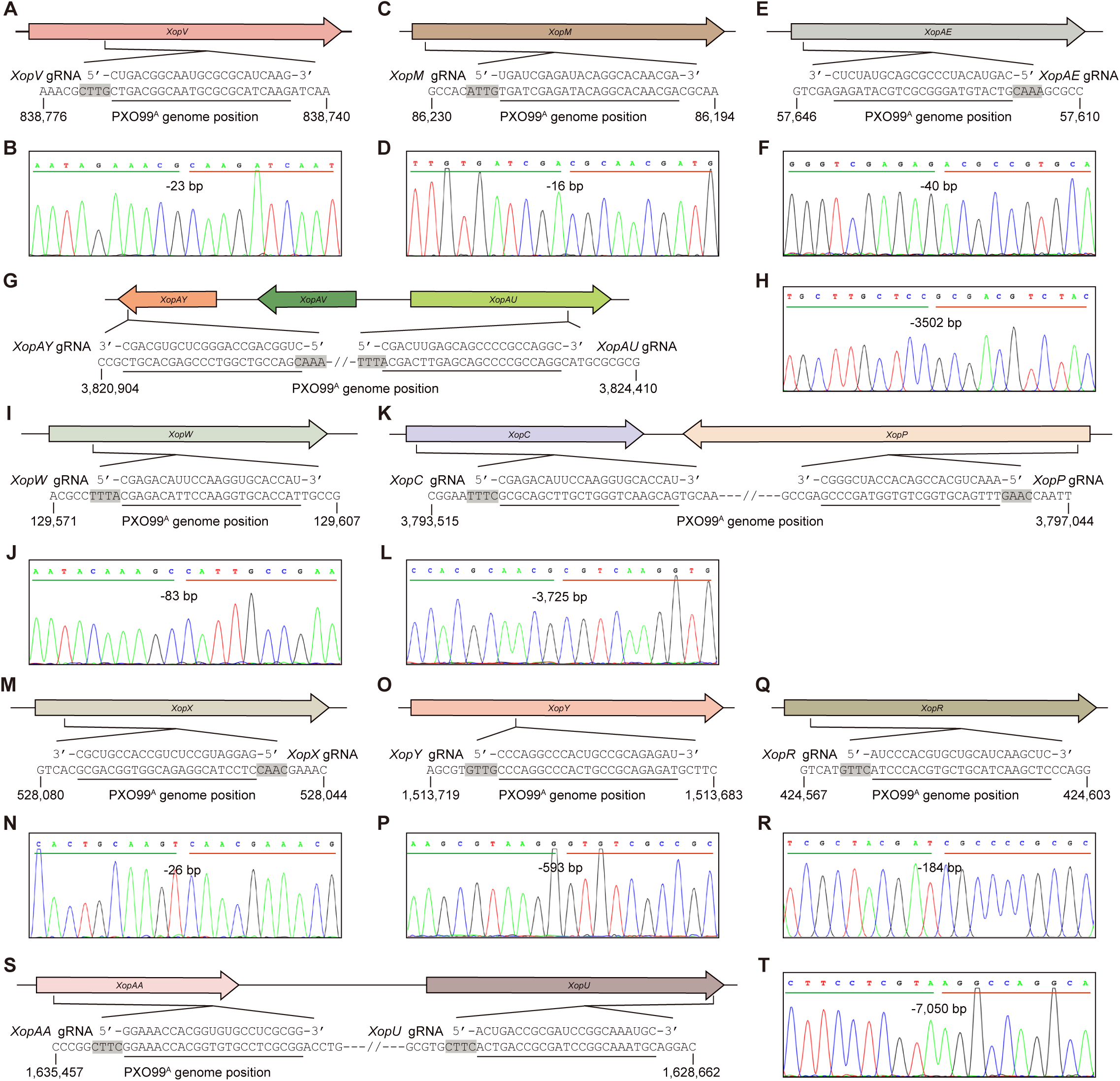
Mutation information on *Xop* genes in the PXO99^A^D25E strain generated by iterative genome editing. Specific gRNAs were designed to target *XopV* **(A)**, *XopM* **(C)**, *XopAE* **(E)**, the *XopAY/AV/AU* gene cluster **(G)**, *XopW* **(I)**, the *XopC/P* gene cluster **(K)**, *XopX* **(M)**, *XopY* **(O)**, *XopR* **(Q)**, and the *XopAA/U* gene cluster **(S)**, respectively. The target regions in the PXO99^A^ genome are underlined and the PAM sequences are marked by the black shadow. The CRISPR/FnCas12a-induced deletions of *XopV* **(B)**, *XopM* **(D)**, *XopAE* **(F)**, the *XopAY/AV/AU* gene cluster **(H)**, *XopW* **(J)**, the *XopC/P* gene cluster **(L)**, *XopX* **(N)**, *XopY* **(P)**, *XopR* **(R)**, and the *XopAA/U* gene cluster **(T)** are shown in Sanger sequencing chromatograms, respectively.

**S4 Fig.**
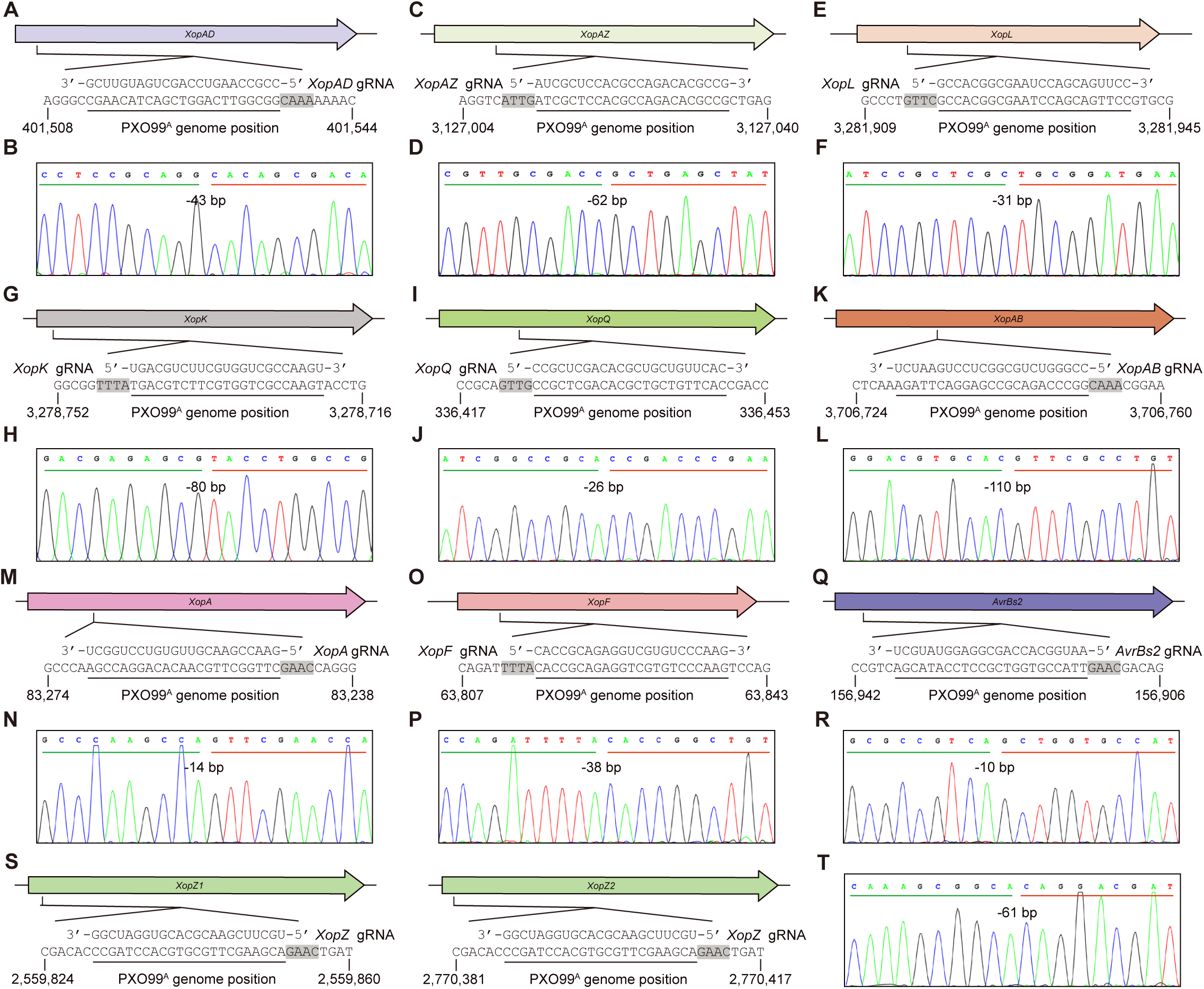
Mutation information on *Xop* genes in the PXO99^A^D25E strain generated by iterative genome editing (continued). Specific gRNAs were designed to target *XopAD* (A), *XopAZ* (C), *XopL* (E), *XopK* (G), *XopQ (I)*, *XopAB* (K), *XopA* (M), *XopF* (O), *AvrBs2* (Q), and *XopZ* (S), respectively. The target regions in the PXO99^A^ genome are underlined and the PAM sequences are marked by the black shadow. Two identical copies of *XopZ* have been annotated in the reference genome of PXO99^A^ (NC_010717.2). The CRISPR/Fn - Cas12a-induced deletions of *XopAD* (B), *XopAZ* (D), *XopL* (F), *XopK* (H), *XopQ* (J), *XopAB* (L), *XopA* (N), *XopF* (P), *AvrBs2* (R), and *XopZ* (T) are shown in Sanger sequencing chromatograms, respectively.

**S5 Fig.**
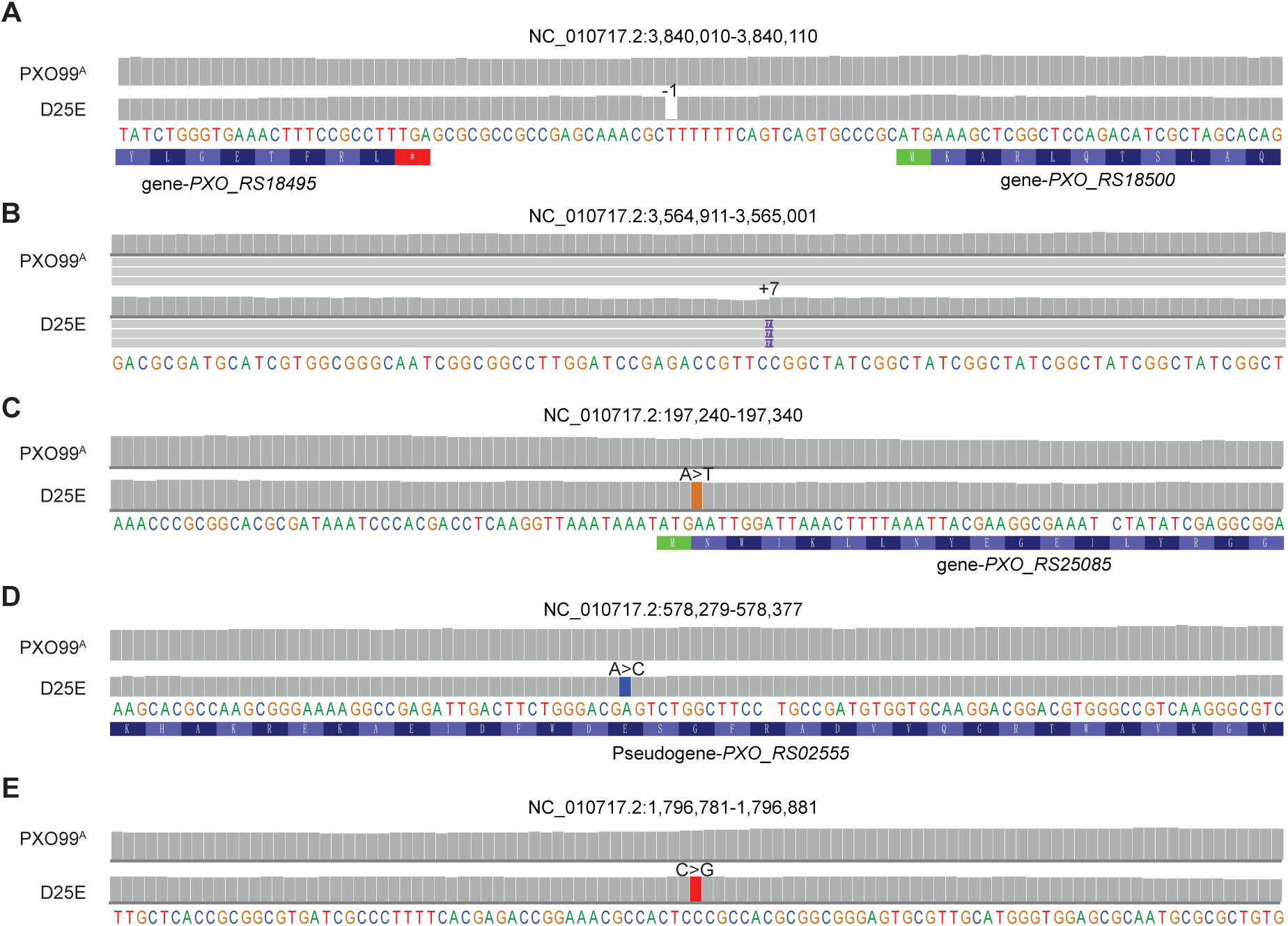
Off-target analysis of *Xoo* PXO99^A^DE25 strain using WGS. IGV genome browser views show 1-bp deletion **(A)**, 7-bp insertion **(B)**, and A-to-G **(C)**, A-to-C **(D)**, and C-to-G **(E)** nucleotide substitutions detected at 5 loci in the PXO99^A^DE25 strain.

**S1 Table.**
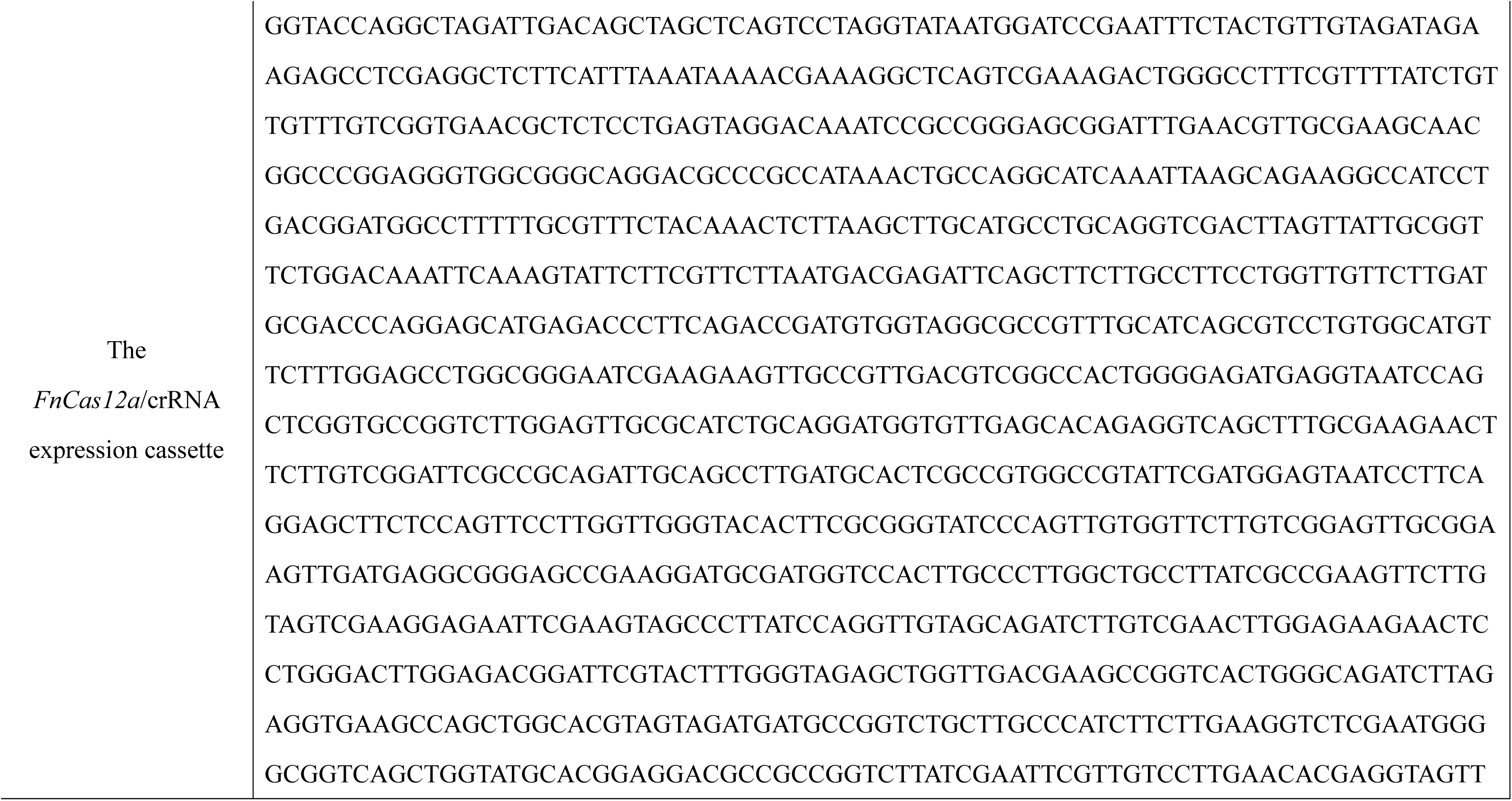
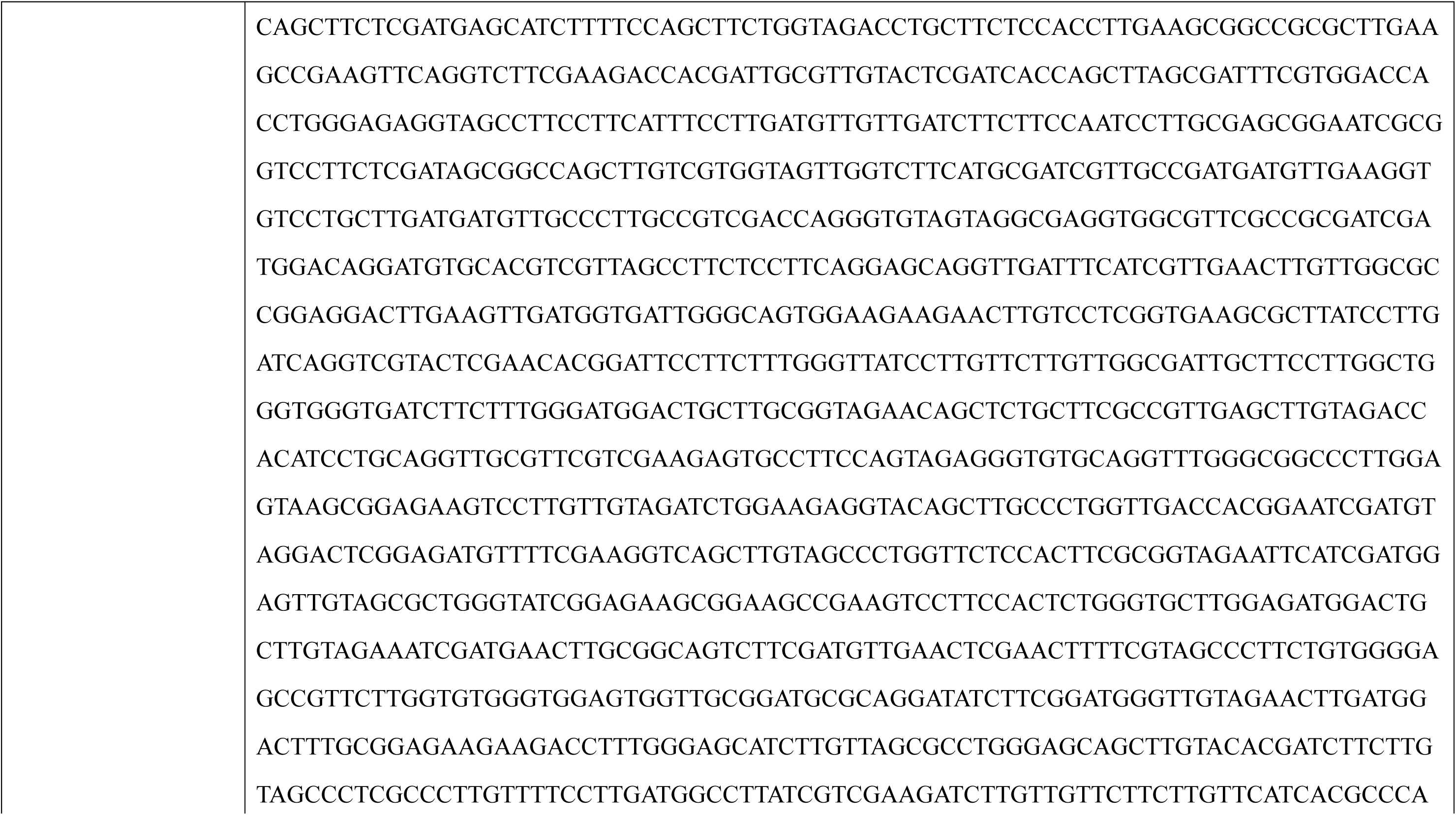
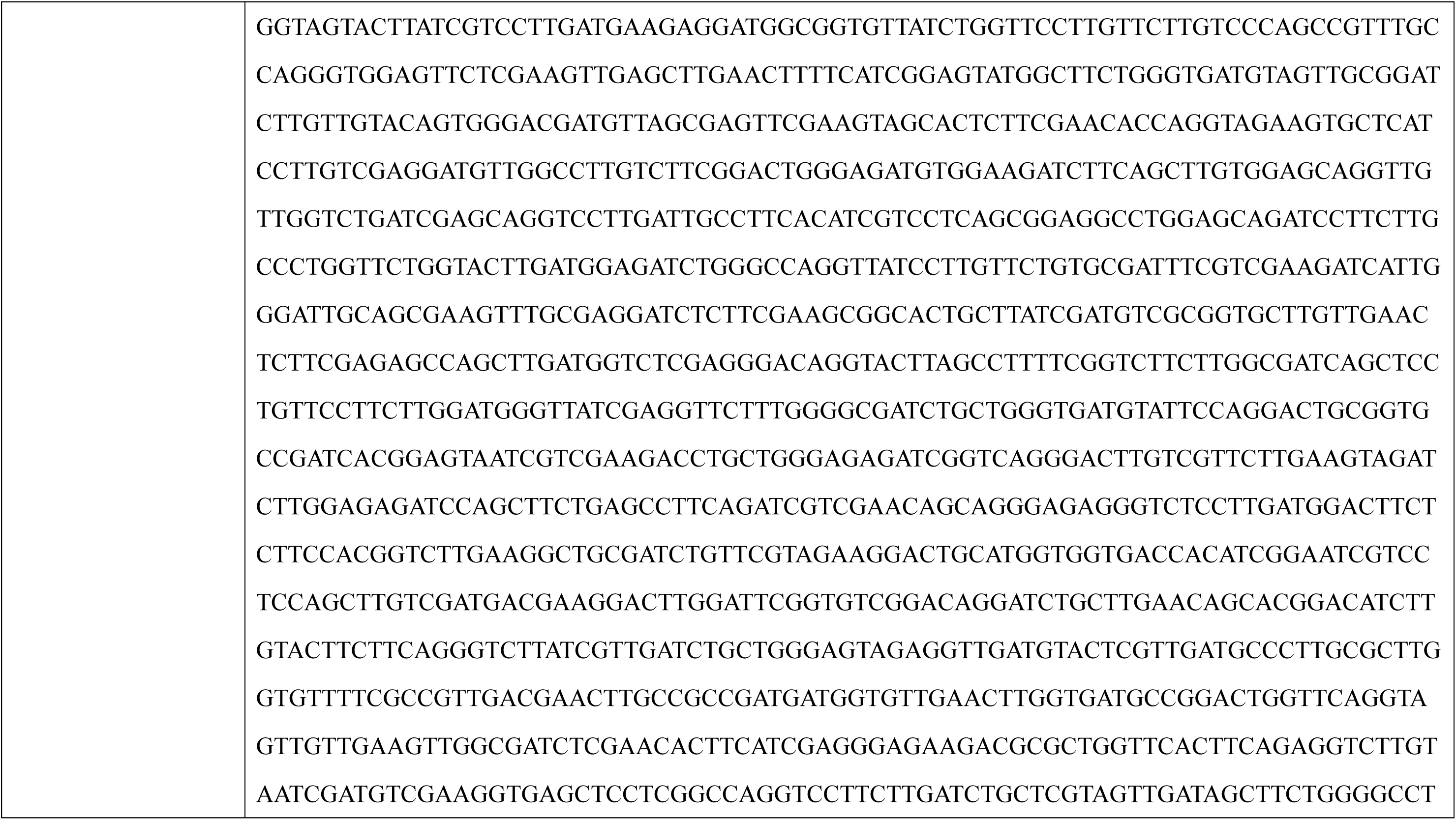
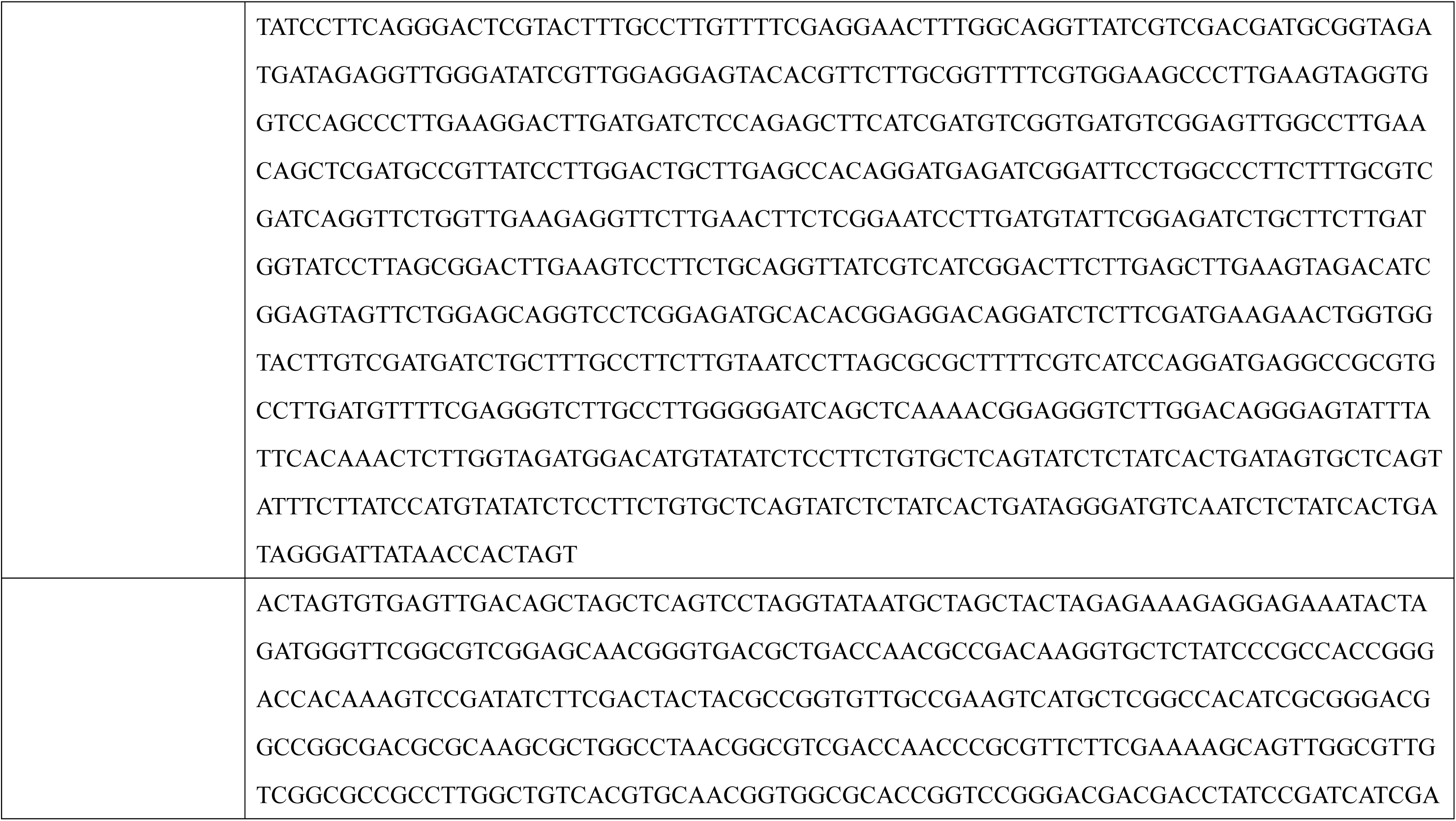
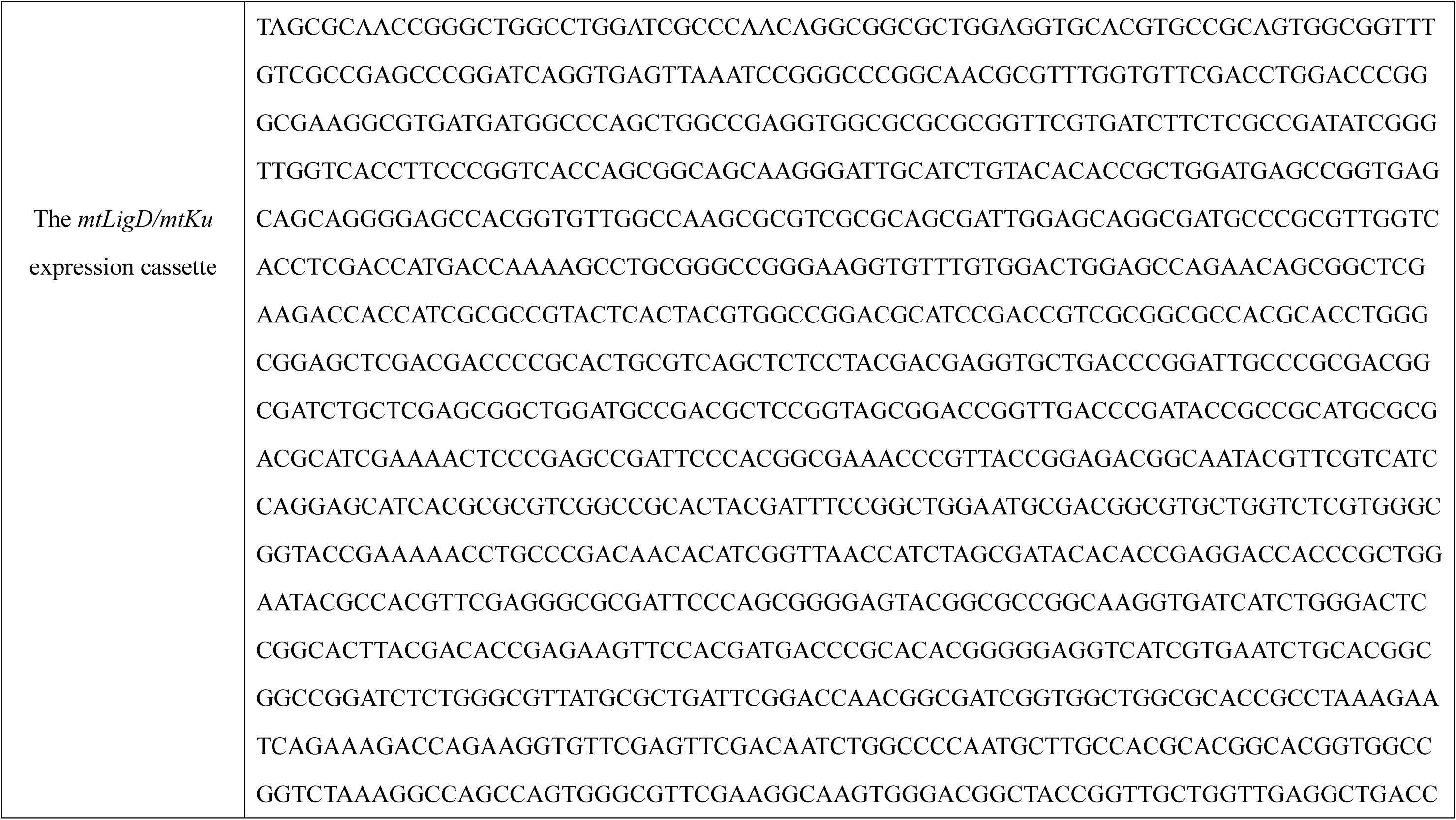
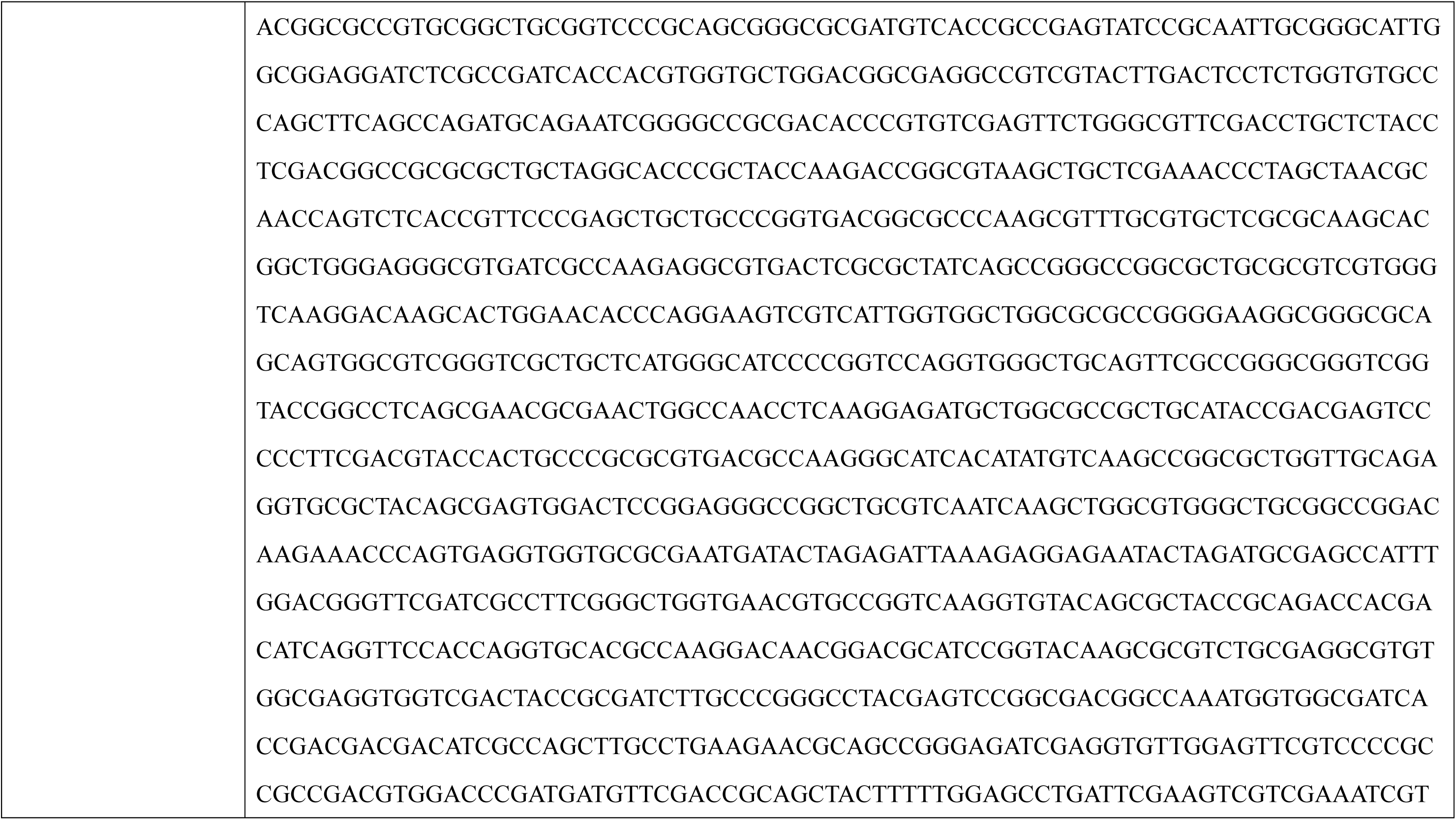
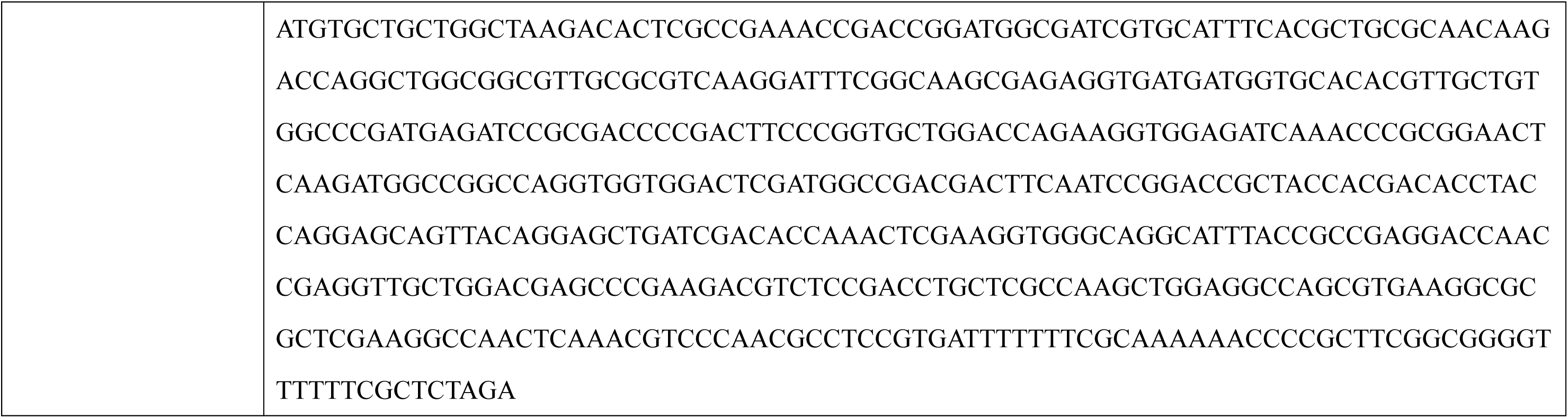
The complete nucleotide sequences of the codon-optimized *FnCas12a* and *mtLigD/mtKu* fragments.

**S2 Table.** Target gene list for genome editing in this study.

| Non-TAL effector genes in *X. oryzae* pv. *oryzae* PXO99<sup>A</sup> |  |  |  |  |  |  |  |
| --- | --- | --- | --- | --- | --- | --- | --- |
|  | Effector Gene | Locus Tag | Genomic Region |  | Effector Gene | Locus Tag | Genomic Region |
| **1** | *XopN* | PXO_RS23030 | 5,033,637..5,035,799 | **14** | *XopF* | PXO_RS00310 | 63,547..65,532 |
| **2** | *XopZ* | PXO_RS11755<br>PXO_RS12685 | 2,559,709..2,563,875<br>2,770,266..2,774,432 | **15** | *XopW* | PXO_RS25045 | 129,539..129,862 |
| **3** | *XopV* | PXO_RS03830 | 838,061..839,056 | **16** | *XopX* | PXO_RS02310 | 526,159..528,267 |
| **4** | *XopK* | PXO_RS14875 | 3,276,325..3,278,856 | **17** | *XopAB* | Annotation missing | 3,706,572..3,707,153 |
| **5** | *XopC* | PXO_RS17330 | 3,793,483..3,794,721 | **18** | *XopAD* | PXO_RS01735 | 401,301..409,760 |
| **6** | *XopP* | PXO_RS17335 | 3,794,949..3,797,090 | **19** | *XopA* | PXO_RS00415 | 82,907..83,326 |
| **7** | *XopQ* | PXO_RS01415 | 336,105..337,499 | **20** | *XopM* | PXO_RS00425 | 84,729..86,282 |
| **8** | *XopY* | Annotation missing | 1,513,135..1,513,980 | **21** | *XopAE* | PXO_RS00290 | 56,059..57,687 |
| **9** | *XopU* | PXO_RS07475 | 1,628,551..1,631,631 | **22** | *XopAV* | PXO_RS17455 | 3,821,959..3,822,735 |
| **10** | *XopAA* | PXO_RS07485 | 1,633,531..1,635,618 | **23** | *XopAY* | PXO_RS17445 | 3,820,875..3,821,630 |
| **11** | *XopR* | PXO_RS24050 | 424,475..425,788 | **24** | *XopAU* | PXO_RS17465 | 3,823,144..3,824,718 |
| **12** | *XopL* | PXO_RS14900 | 3,281,677..3,283,635 | **25** | *XopAZ* | PXO_RS14270 | 3,126,943..3,127,416 |
| **13** | *AvrBs2* | PXO_RS00725 | 154,940..157,084 |  |  |  |  |

| Genes in *X. oryzae* pv. *oryzae* PXO99<sup>A</sup> |  |  |  |  |  |  |  |
| --- | --- | --- | --- | --- | --- | --- | --- |
|  | Gene Name | Locus Tag | Genomic Region |  | Gene Name | Locus Tag | Genomic Region |
| 1 | - | PXO_RS06855 | 1,475,772..1,480,253 | 2 | - | PXO_RS06285 | 1,349,572..1,354,062 |
| Effector genes in *P. syringae* pv. *tomato* DC3000 |  |  |  |  |  |  |  |
|  | Effector Gene | Locus Tag | Genomic Region |  | Effector Gene | Locus Tag | Location |
| 1 | *HopAB2* | PSPTO_3087 | 3,468,470..3,470,131 | 2 | *HopX1* | PSPTO_A0012 | 14,914..16,068 |

**S3 Table.**
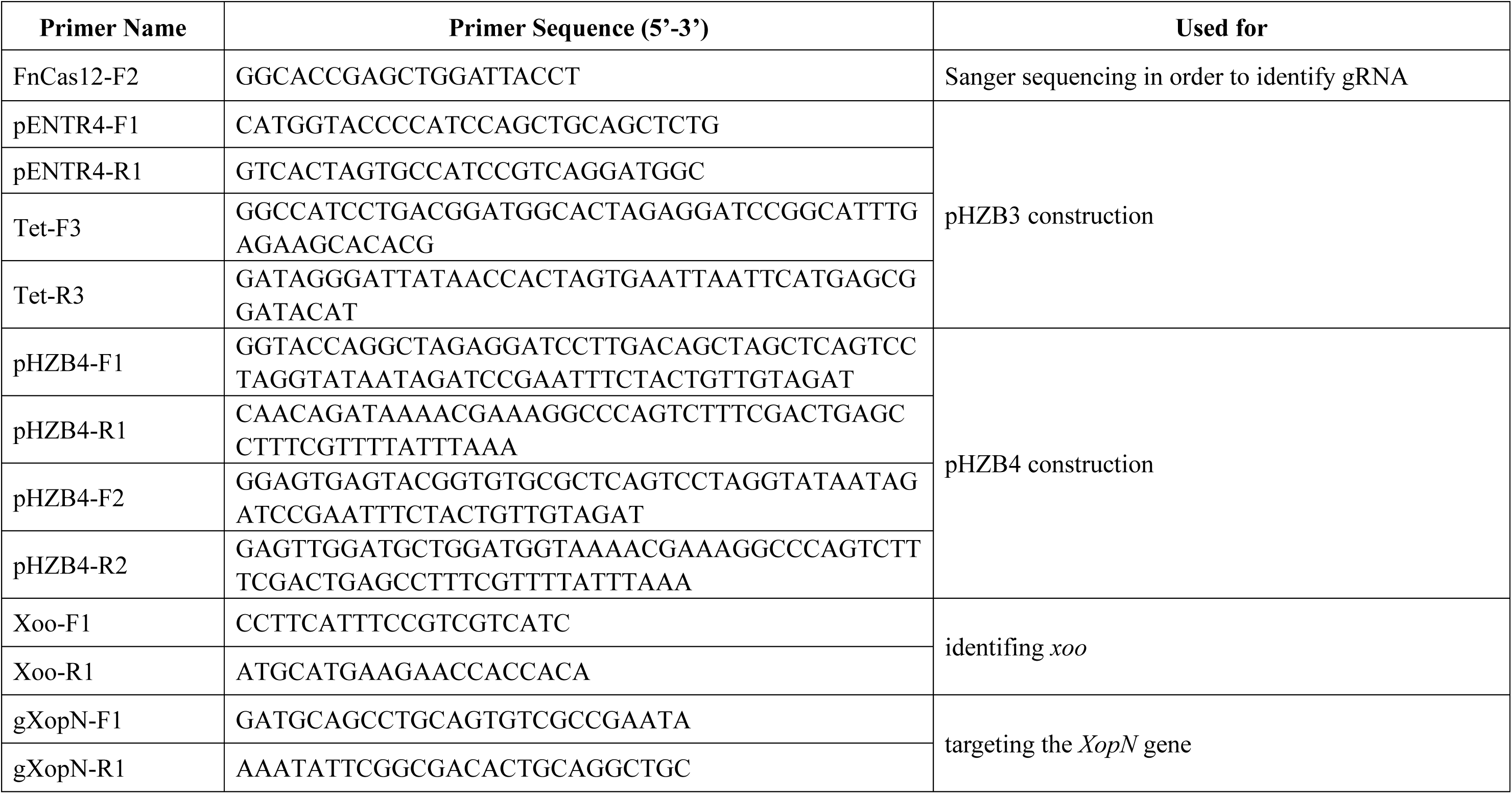
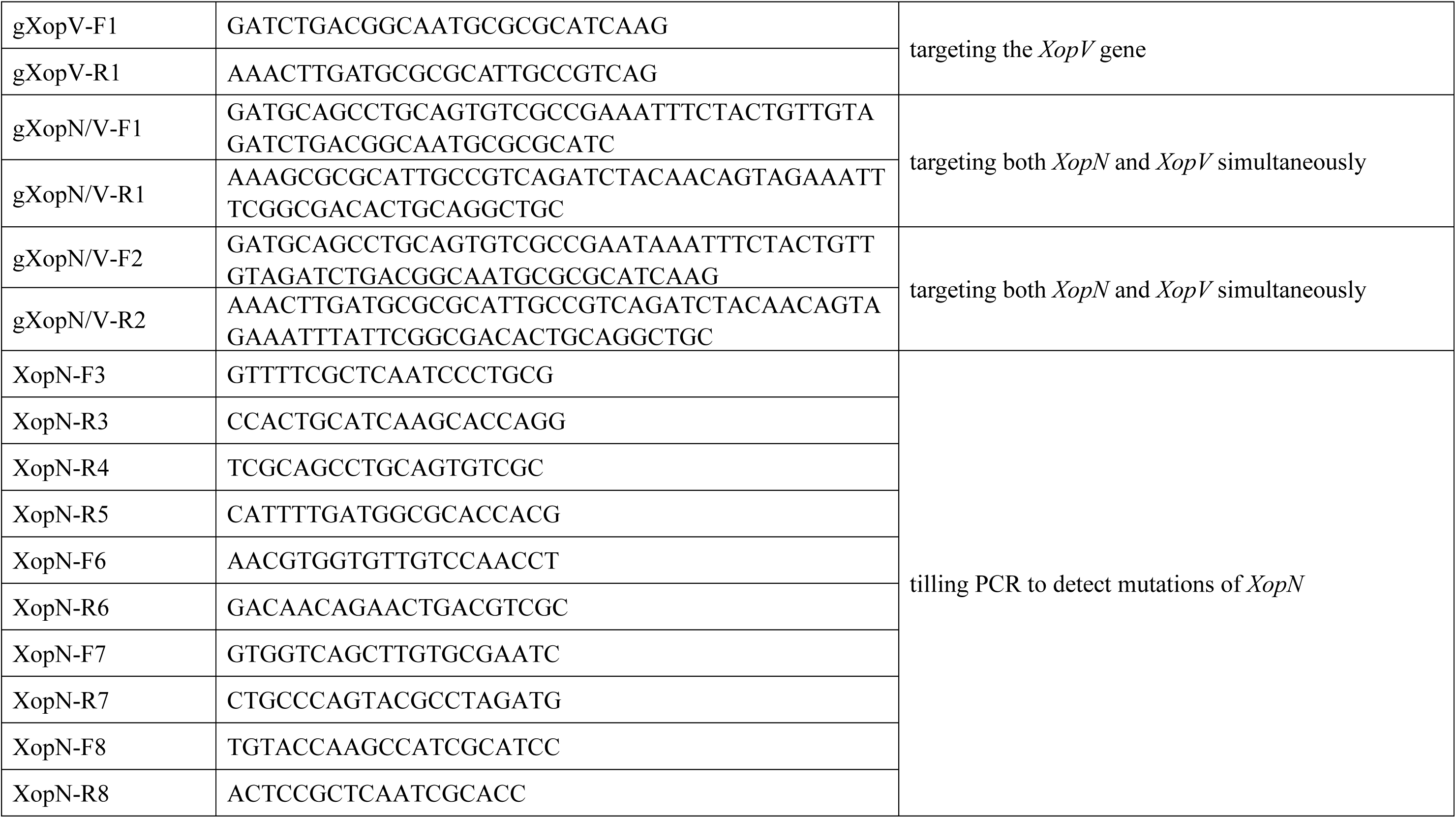
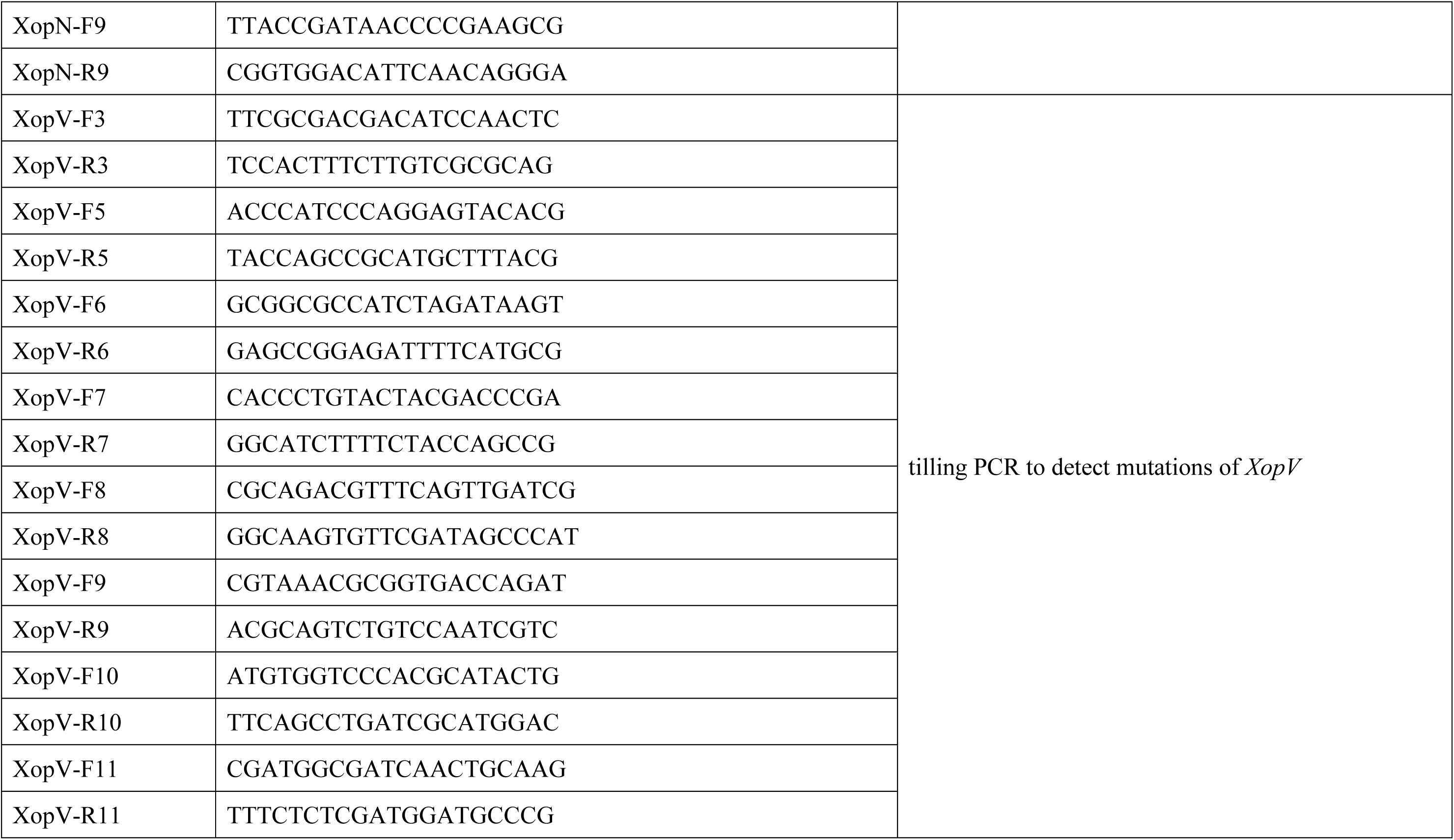
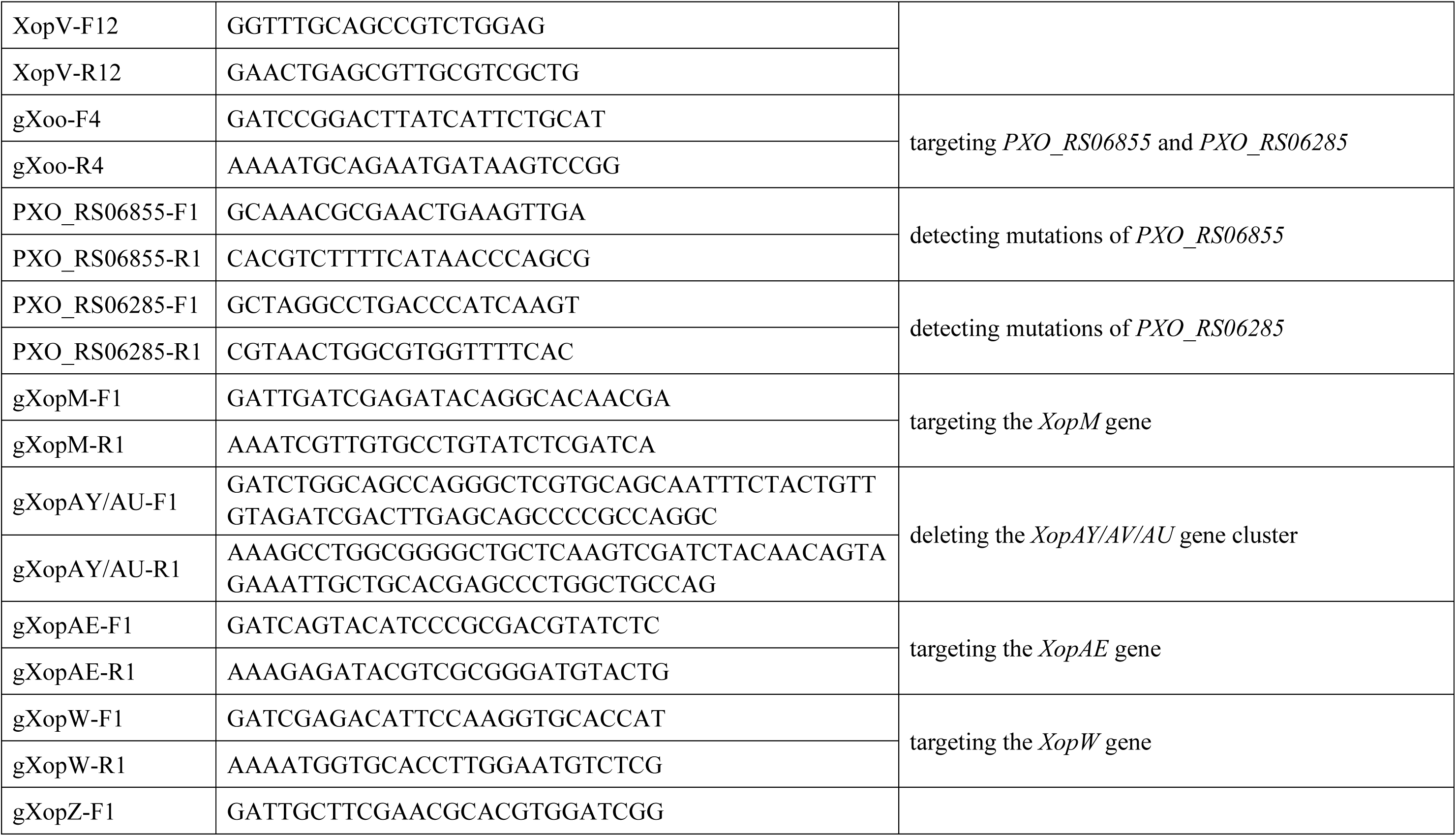
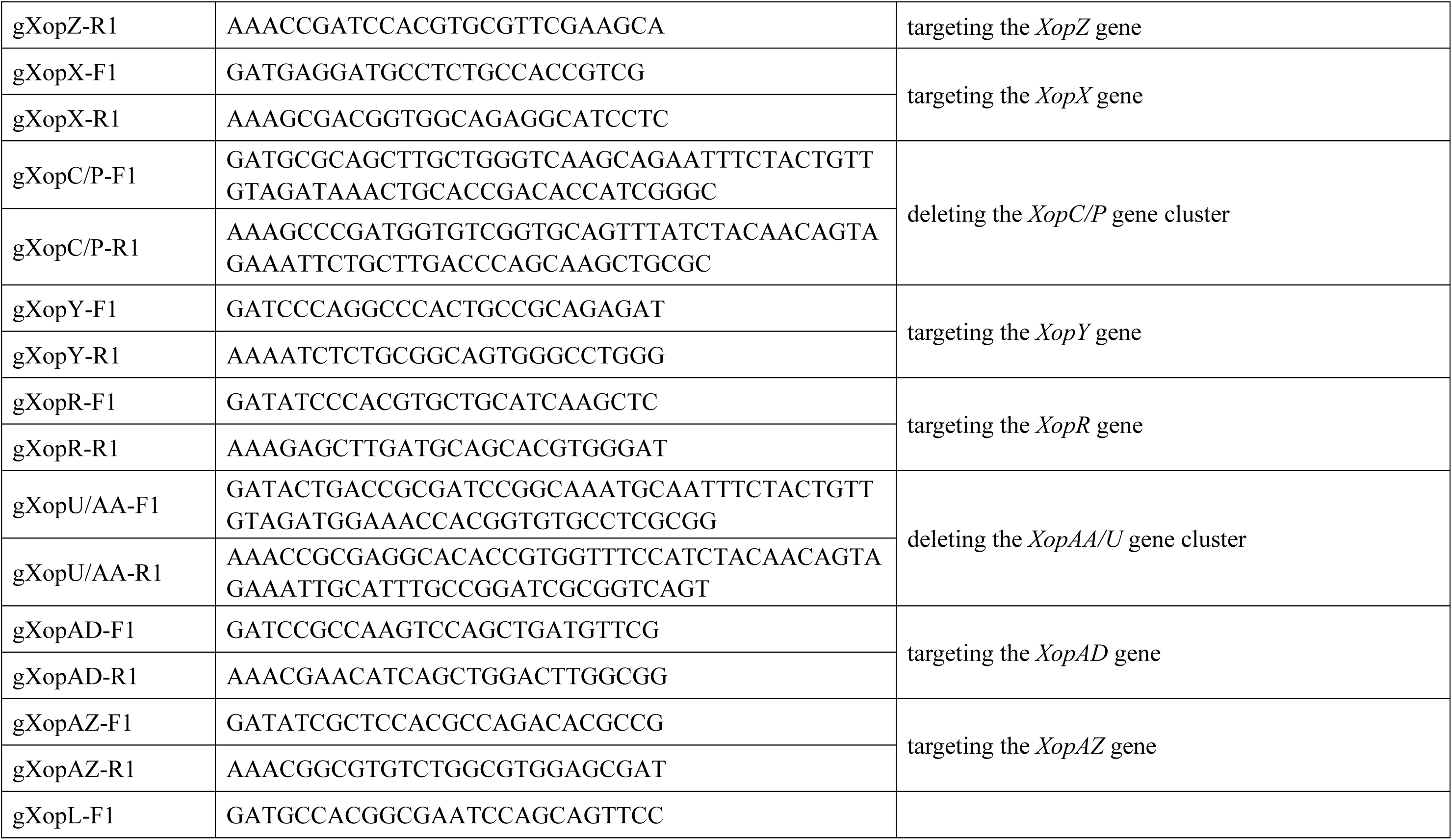
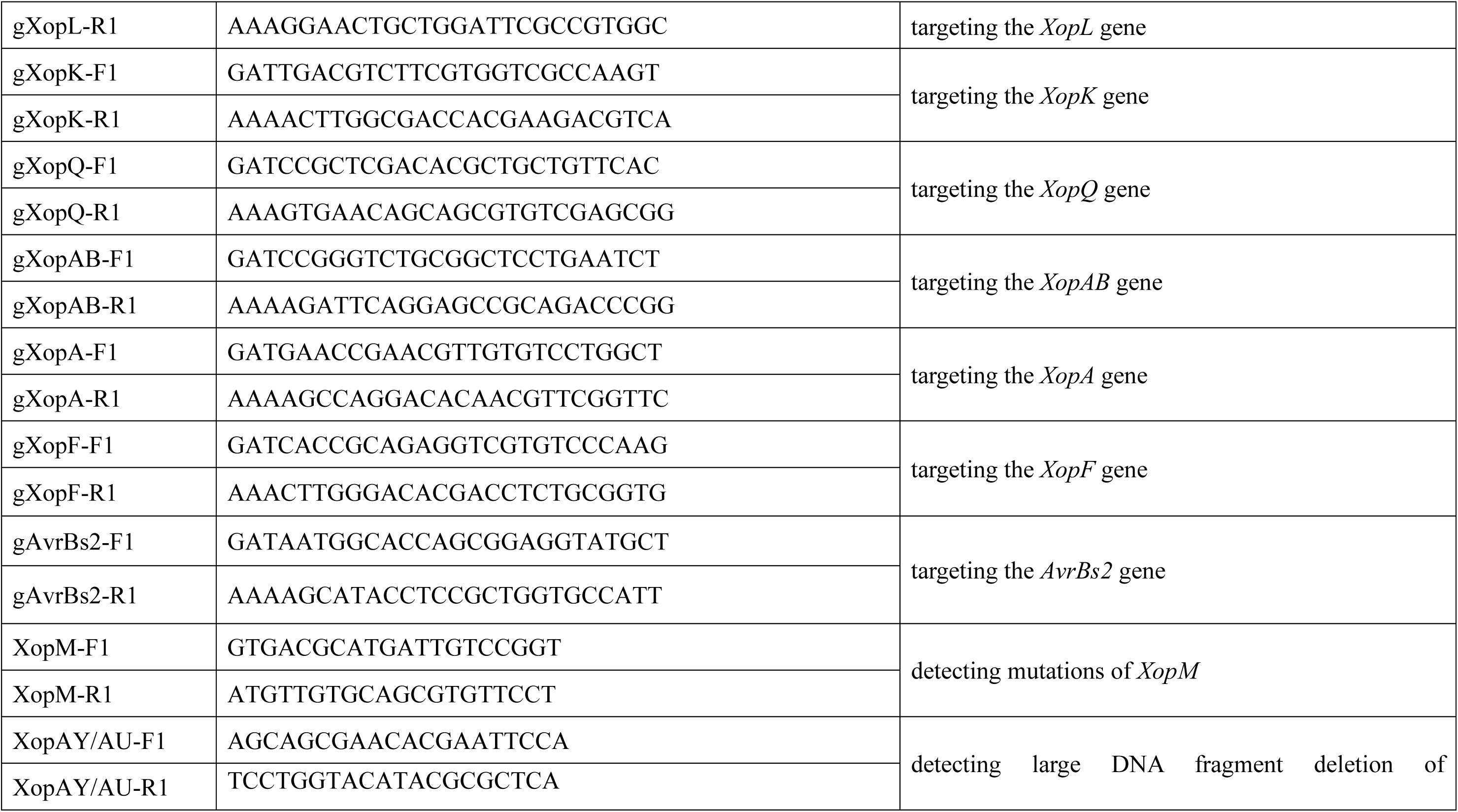
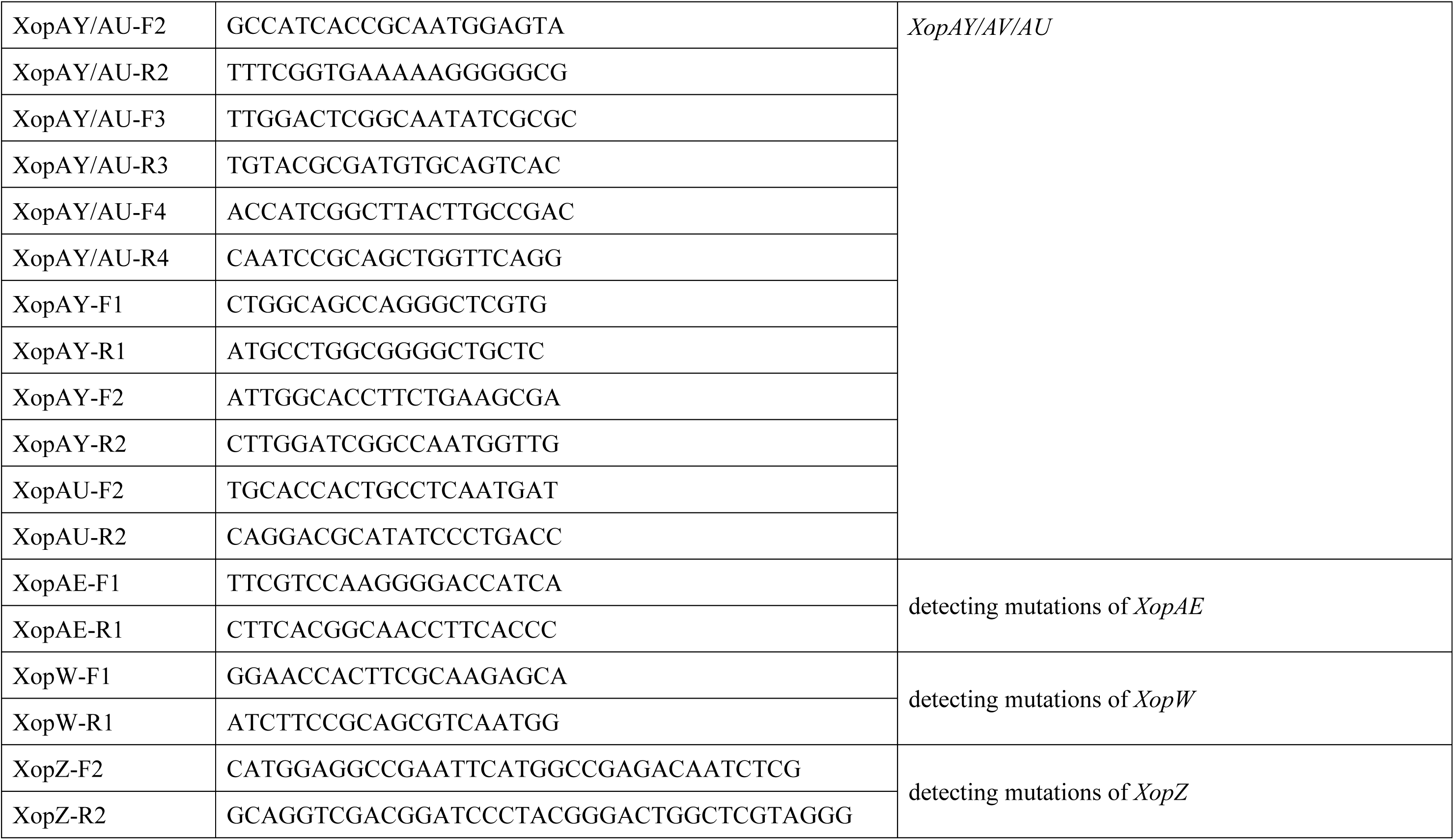
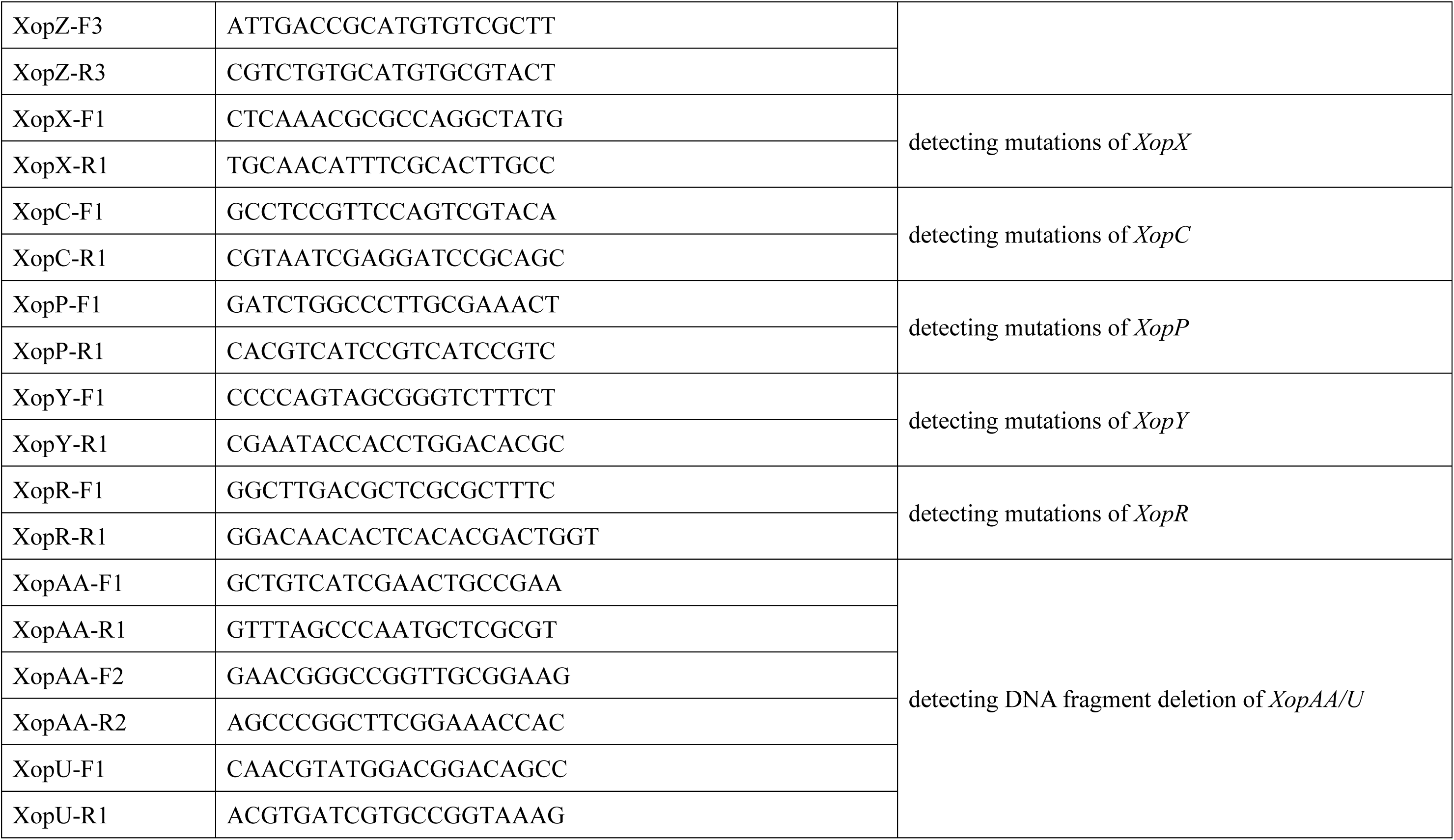
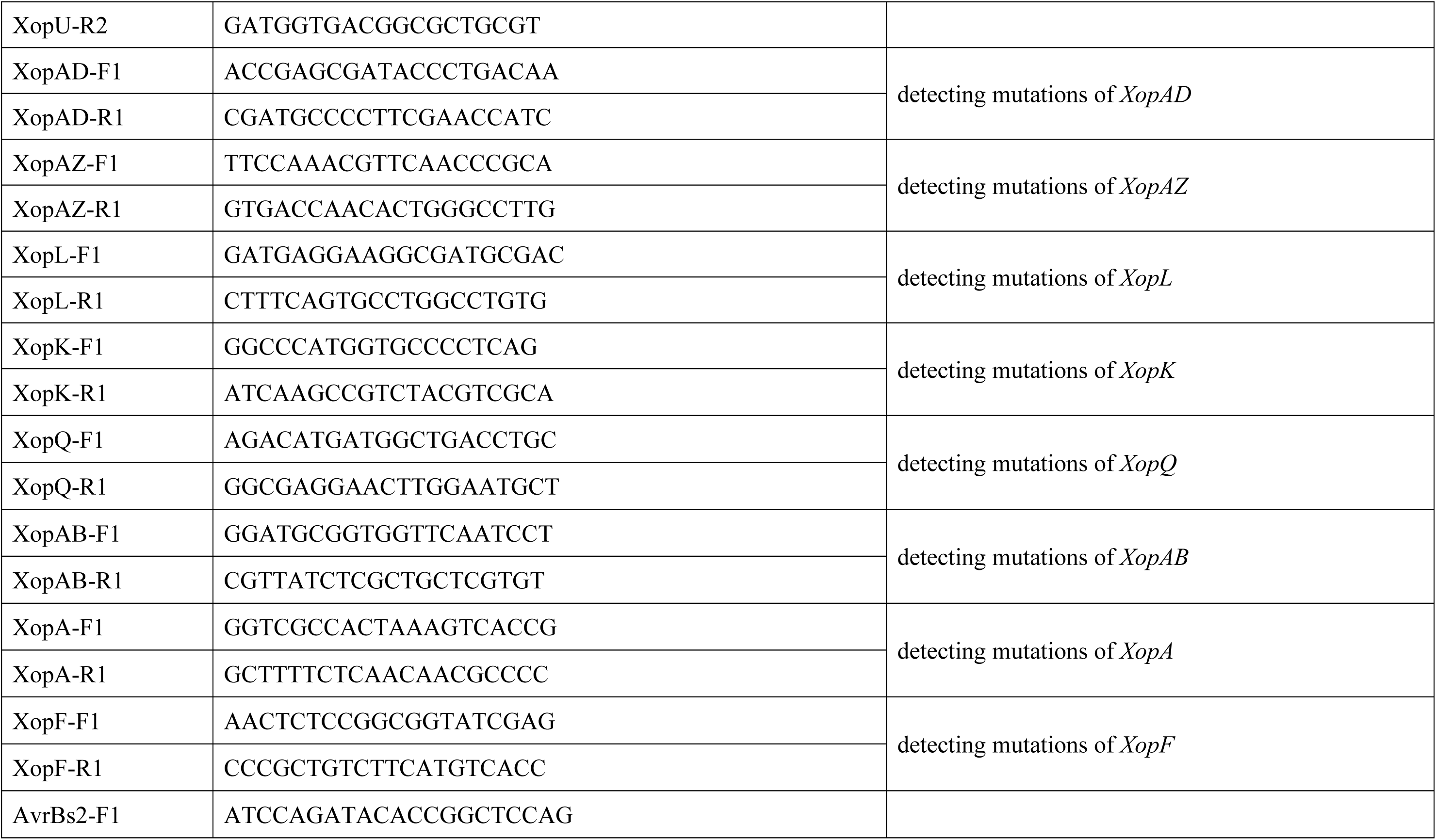
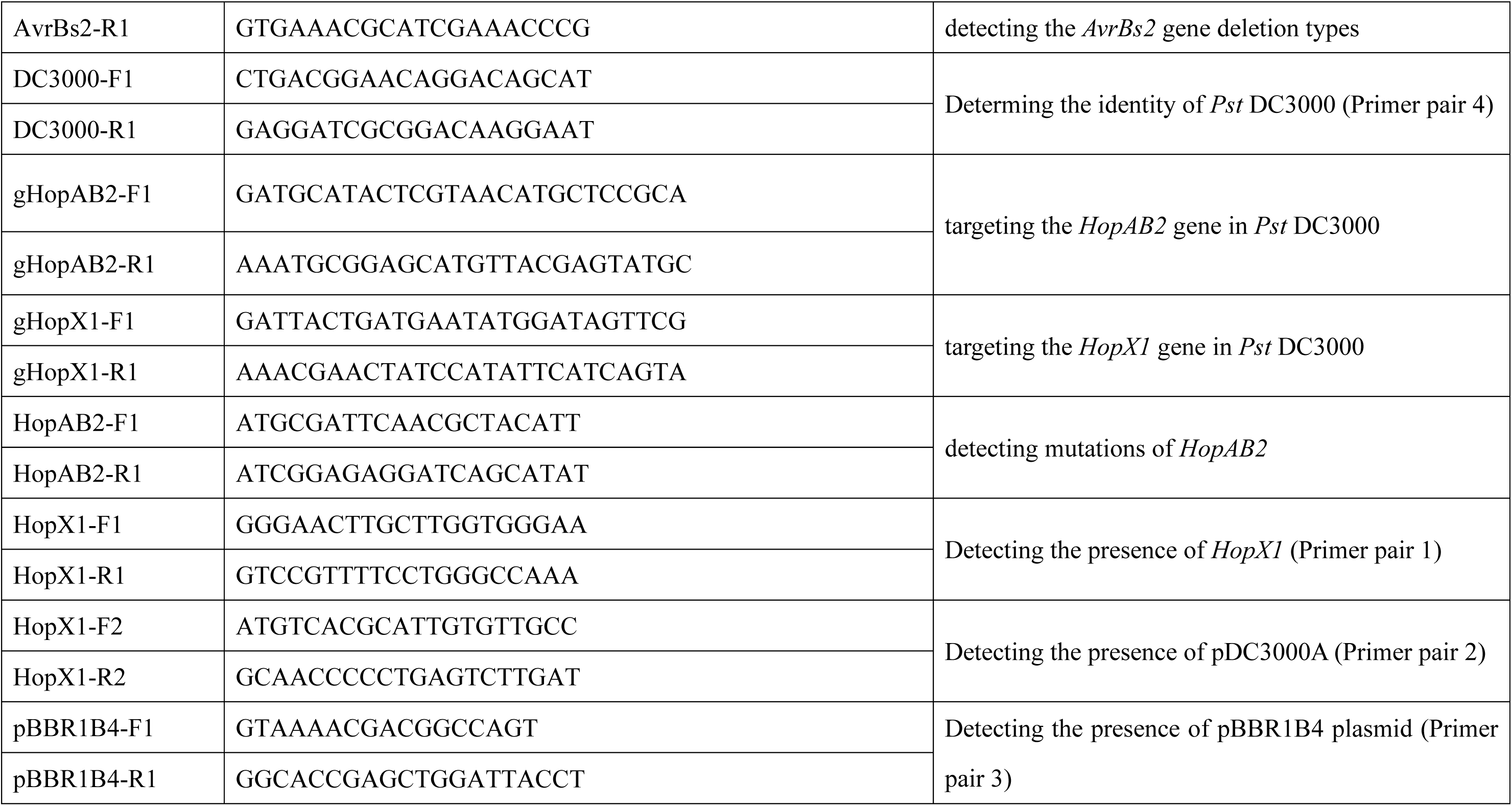
Oligonucleotides used in this study.

**S4 Table.**
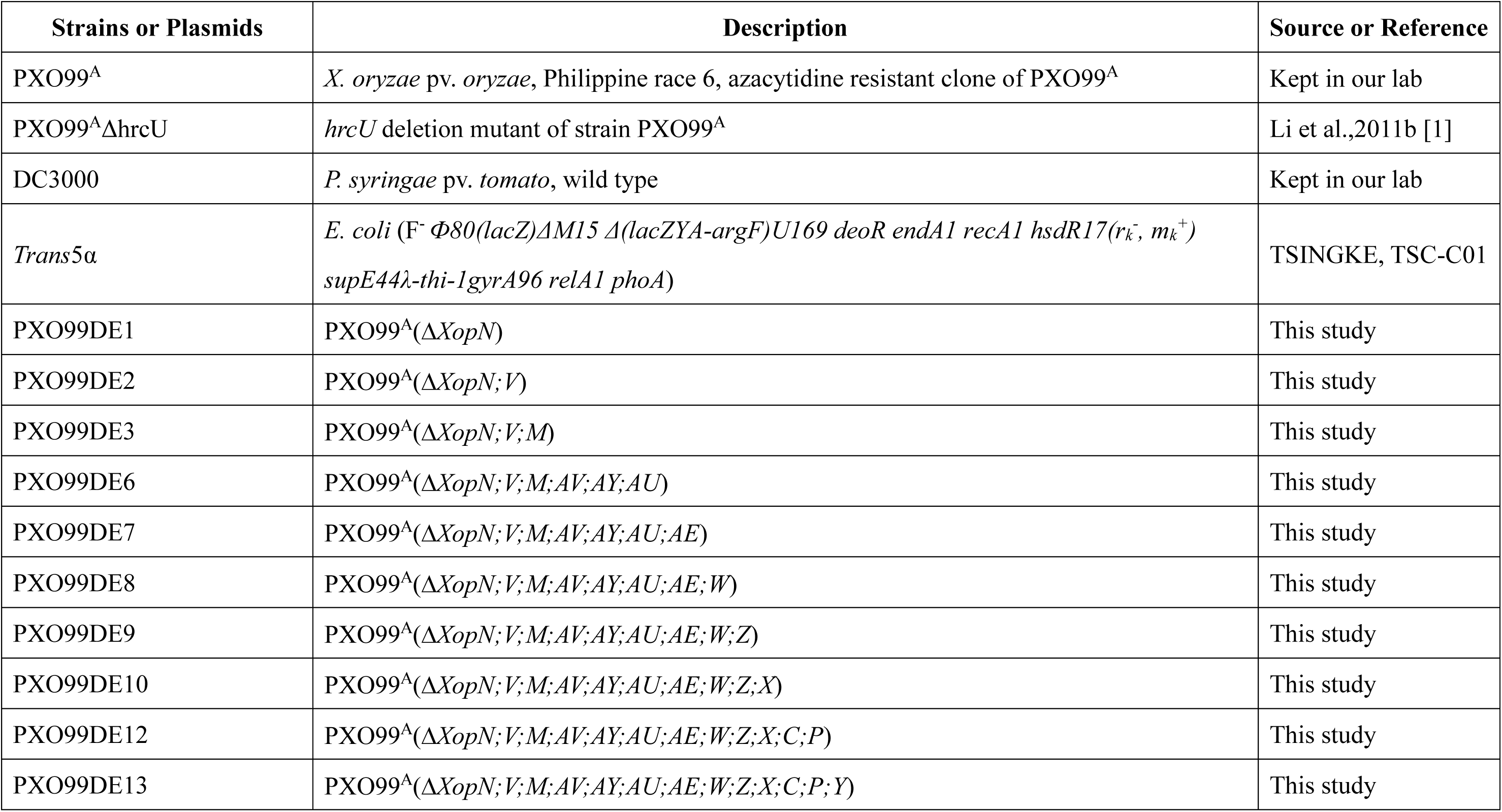
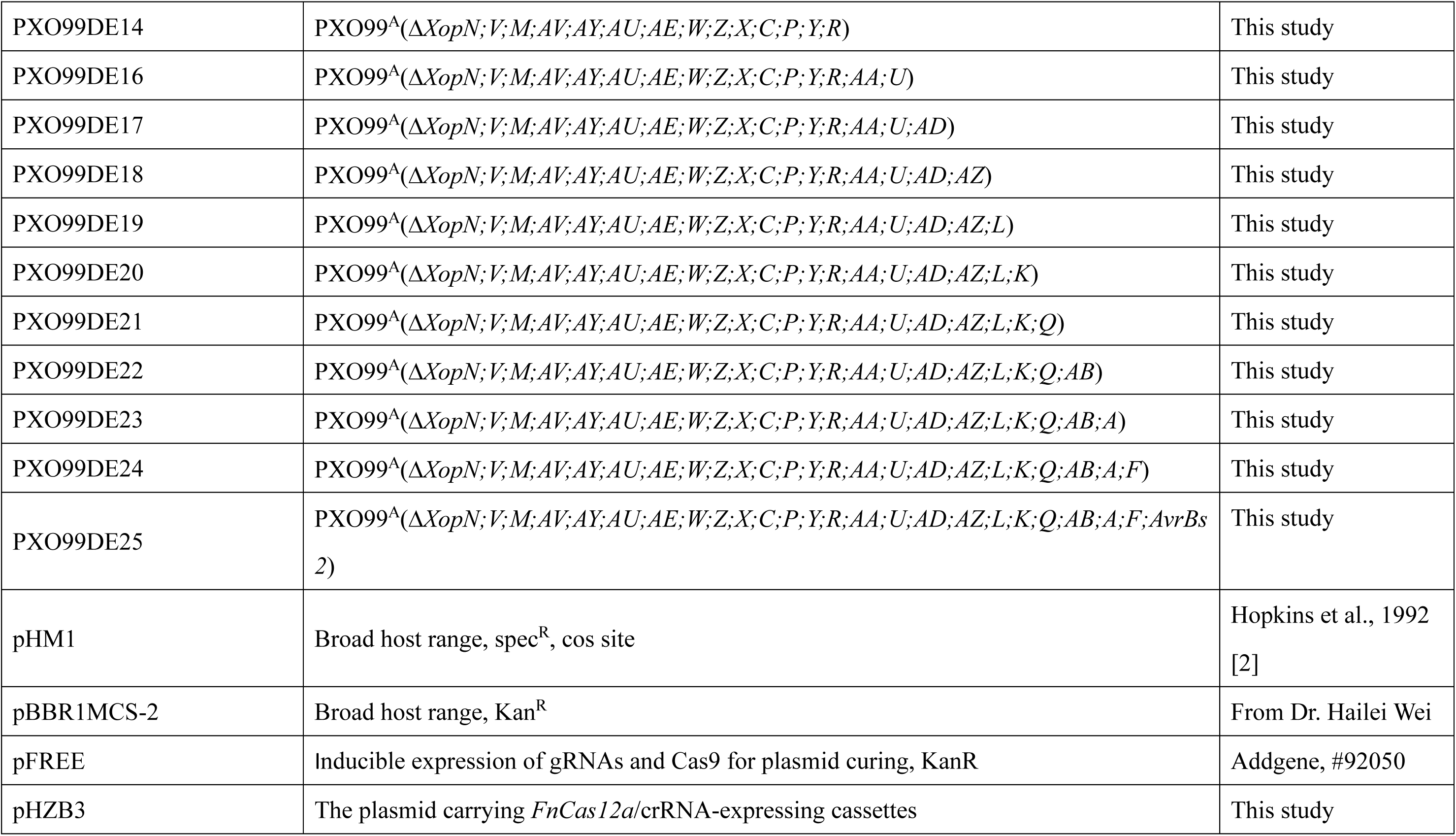
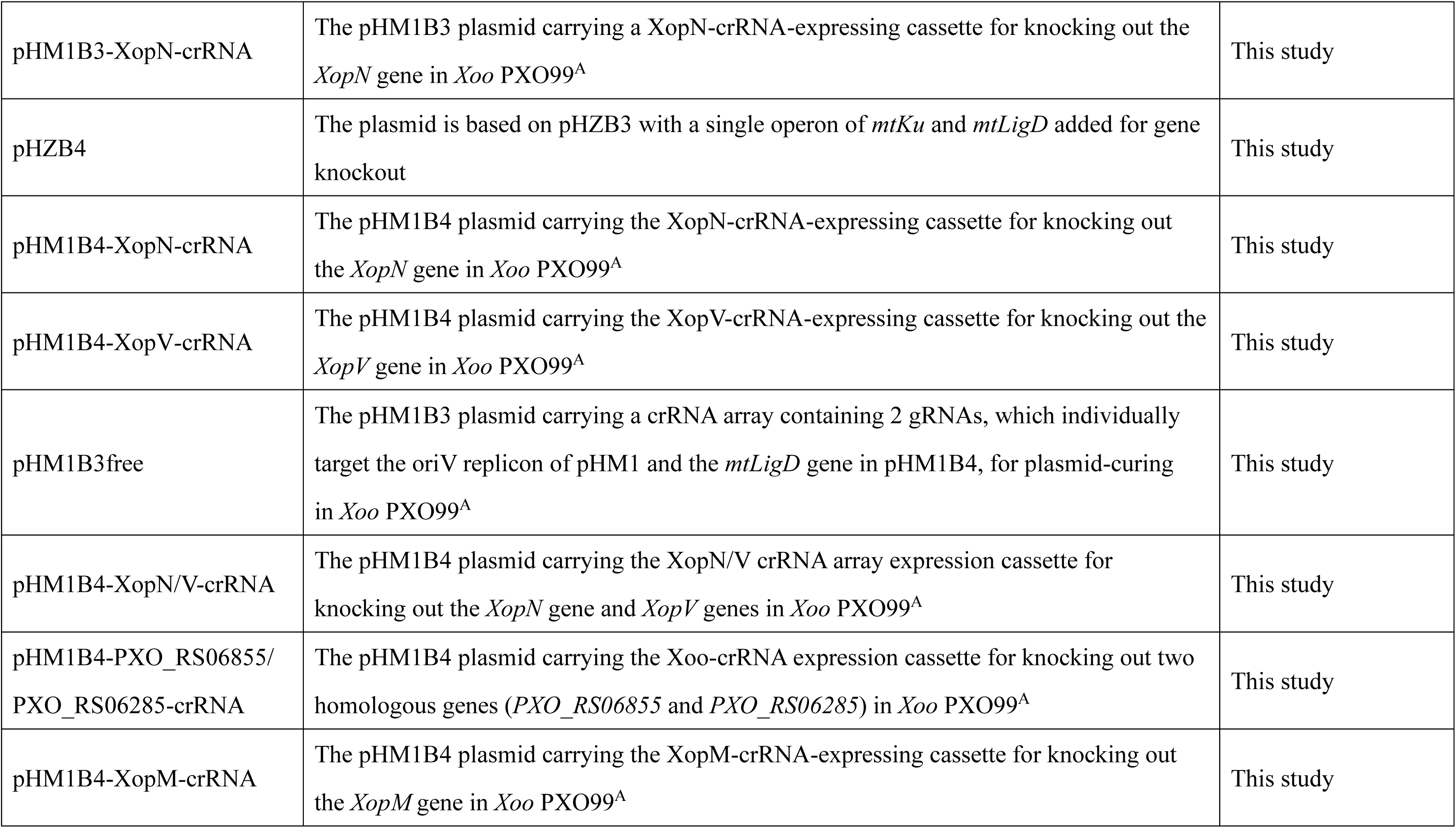
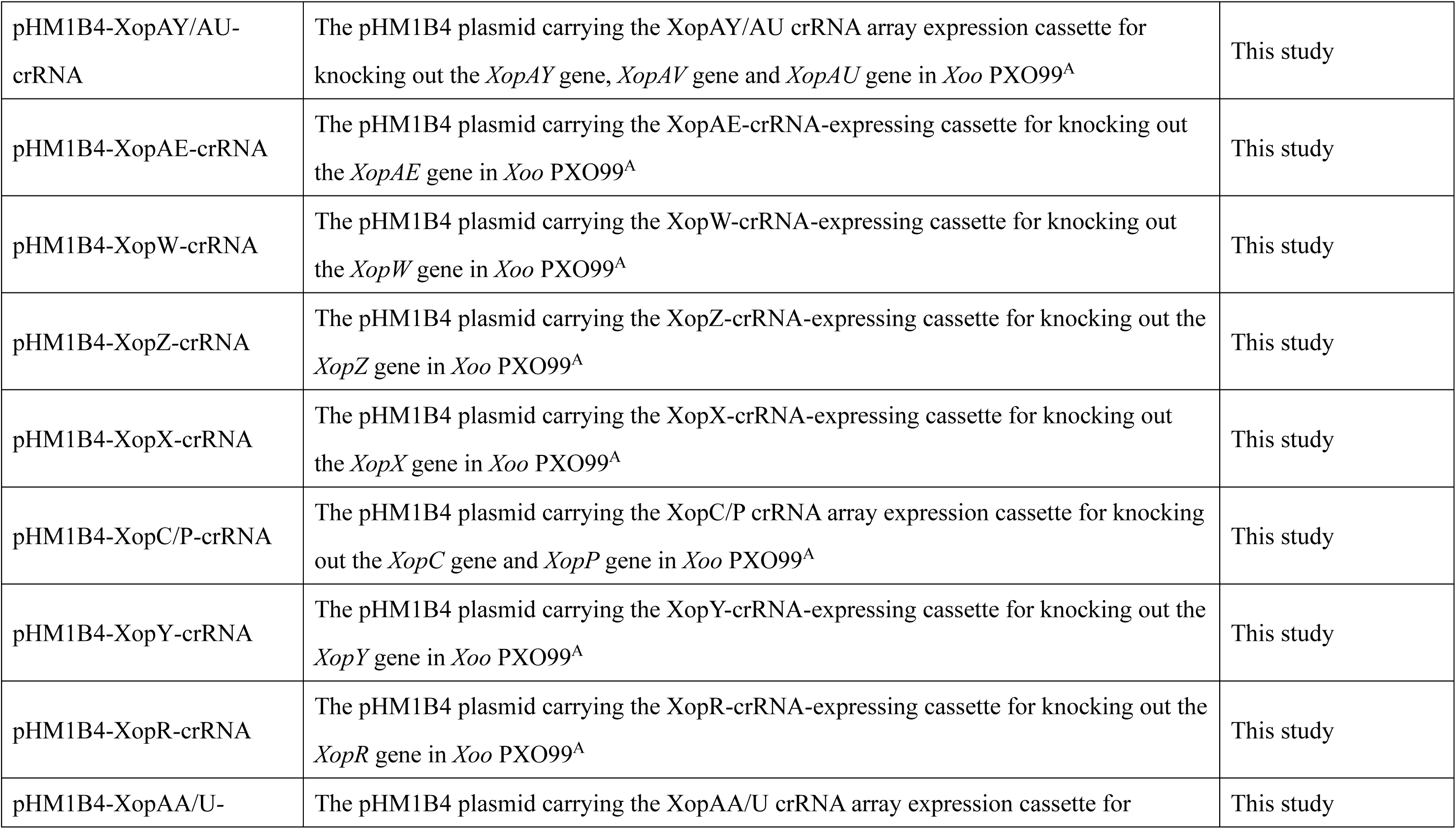
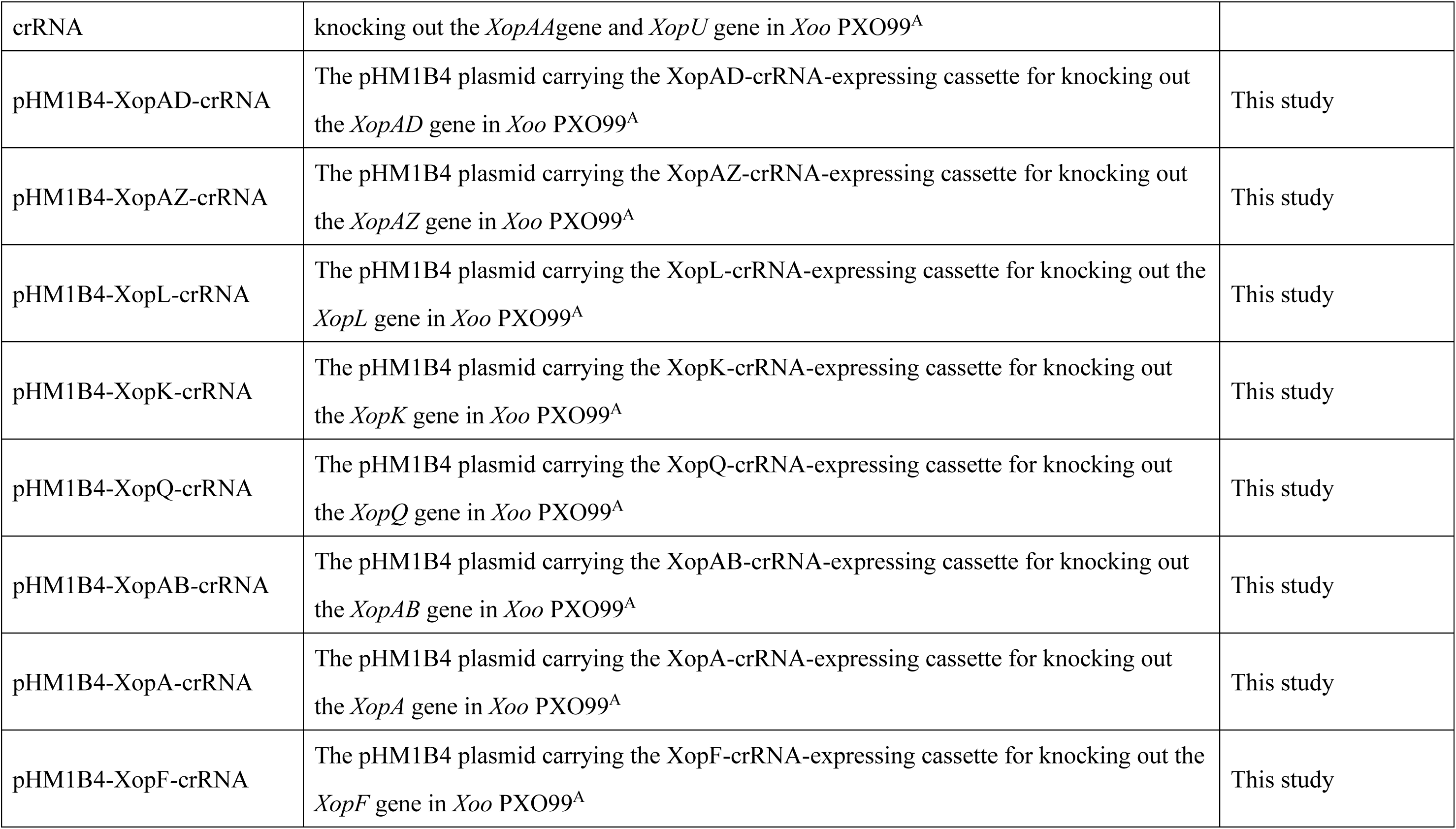
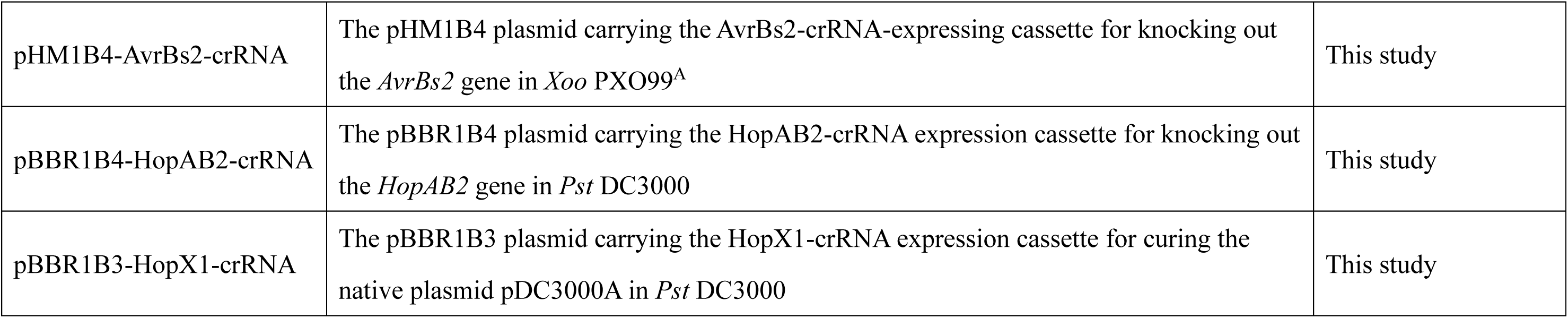
Bacterial strains and plasmids used in this study.

